# ABIN1 is a negative regulator of effector functions in cytotoxic T cells

**DOI:** 10.1101/2023.10.25.563918

**Authors:** Sarka Janusova, Darina Paprckova, Juraj Michalik, Valeria Uleri, Ales Drobek, Eva Salyova, Louise Chorfi, Ales Neuwirth, Jan Prochazka, Radislav Sedlacek, Peter Draber, Ondrej Stepanek

**Author notes:** The correspondence should be addressed to Ondrej Stepanek.

## Abstract

T cells are pivotal in the adaptive immune defense, necessitating a delicate balance between robust response against infections and self-tolerance. Their activation involves intricate cross-talk among signaling pathways triggered by the T-cell antigen receptors (TCR) and co-stimulatory or inhibitory receptors. The molecular regulation of these complex signaling networks is still incompletely understood. We identified an adaptor protein, ABIN1 as a component of the signaling complexes of GITR and OX40 co-stimulation receptors. T cells lacking ABIN1 are hyper-responsive ex vivo, and exhibit enhanced responses to cognate infections, and superior ability to induce experimental autoimmune diabetes in mice. We observed that ABIN1 negatively regulates NF-κB and p38 pathways. The latter was at least partially responsible for the upregulation of key effector proteins, IFNG and GZMB in ABIN1-deficient T cells after TCR stimulation. Our findings reveal the intricate role of ABIN1 in T-cell regulation and its potential as a target for therapeutic fine-tuning of T-cell responses.

## Introduction

T cells use their variable T-cell antigen receptors (TCR) to recognize cognate antigens to initiate adaptive immune responses to infections and cancer. To prevent overt T-cell responses which might lead to autoimmune pathology, a plethora of mechanisms regulating T-cell activation has evolved, including the co-stimulatory and inhibitory surface receptors and intracellular regulators of signaling pathways. Understanding the individual regulatory steps of T-cell activation is critical for developing novel immunotherapeutic strategies to treat T cell-related diseases such as cancer, infection, and autoimmunity. Several co-stimulatory receptors from the TNFR superfamily (e.g., GITR, OX40, CD137) are up-regulated upon the initial antigen encounter. As these receptors enhance T-cell responses [1], some of them are used as targets of emerging anti-tumor immunotherapies [2–5]. However, despite the recent progress in uncovering the CD137 signalosome [6], the mechanisms of signal transduction of these co-stimulation TNFR superfamily receptors are still incompletely understood.

A20-binding inhibitor of NFκB 1 (ABIN1 alias TNIP1) is an adaptor protein interacting with polyubiquitin signaling chains, NF-kappa-B essential modulator (NEMO), and a deubiquitinase, A20 [7]. It has been shown that ABIN1 negatively regulates several signaling pathways, which use polyubiquitin chains for signal propagation, such as tumor-necrosis factor receptor 1 (TNFR1), toll-like receptors, and autophagy signaling [7–12]. Polymorphisms in *ABIN1* are associated with systemic lupus erythematosus in humans [13]. Mice deficient in *Abin1* and mice expressing a severely compromised variant of *Abin1* develop a profound phenotype of either TNF-triggered embryonal lethality [8, 14] or a severe autoimmunity [11], depending on the character of the *Abin1* modification and/or the genetic background of the mice. Whereas the importance of ABIN1 for the homeostasis and tolerance of both myeloid and B cells is well documented [10, 11], much less is known about the role of ABIN1 in T cells. A recent study identified ABIN1 as a component of the CARD11-BCL10-MALT1 (CBM) complex [15], which is formed upon TCR signaling in T cells. Subsequent examination of the role of ABIN1 in a T-cell line Jurkat indicated its role in regulating the TCR-induced NF-κB pathway [15]. However, the role of ABIN1 in the biology of primary T cells has not been elucidated.

In this study, we characterized ABIN1 as a component of proximal signaling complexes of co-stimulatory receptors GITR and OX40, suggesting that ABIN1 negatively regulates both the antigenic signaling and T-cell co-stimulation. Using ABIN1-deficient primary T cells, we showed that ABIN1 limits the effector responses of cytotoxic T cells during infection and autoimmunity.

## Results

### ABIN1 is a component of the proximal signaling complexes of GITR and OX40 receptors

Because the signal transduction pathway of co-stimulatory T-cell receptors GITR and OX40 was not completely understood, we analyzed the composition of GITR and OX40 proximal signaling complexes (SC) using a proteomic approach previously used for the characterization of the TNFR1 and IL-17R SCs [16–18]. First, we produced a tagged recombinant GITRL, which binds to PMA/ionomycin-preactivated T cells (Fig. EV1A). Next, we used the GITRL to trigger the GITR signaling pathway in these cells and followed with the cell lysis, purification of the ligand-receptor-SC by tandem affinity purification, and finally by mass spectrometry analysis to identify the SC composition (Fig. 1A, Fig. EV1B-C). The samples in which the ligand was added post-lysis, served as negative controls to reveal contaminating proteins (Fig. EV1B-C). We identified known members of the GITR-SC including TRAF1, TRAF2, CIAP2 [1] together with novel components, namely NF-kappa-B essential modulator (NEMO); subunits of the linear ubiquitin chain assembly complex (LUBAC): HOIP, HOIL1 and SHARPIN; A20 deubiquitinase; and adaptor protein ABIN1 (Fig. 1B-C). We confirmed the recruitment of ABIN1 and A20 into the GITR-SC by immunoprecipitation in DO11.10 hybridoma cells (Fig. 1D) and in primary murine OT-I *Rag2*^KO/KO^ T cells (Fig. 1E). In the next step, we applied the same approach to another co-stimulatory TNFR superfamily receptor, OX40, to identify ABIN1 and A20 as novel components of the OX40-SC (Fig. EV1D-G). Overall, these experiments showed that A20 and ABIN1 are parts of the SC of GITR and OX40, and likely other co-stimulatory T-cell receptors from the TNFR superfamily and are thus, candidate negative regulators of T-cell activation through co-stimulation.

**Figure 1.**
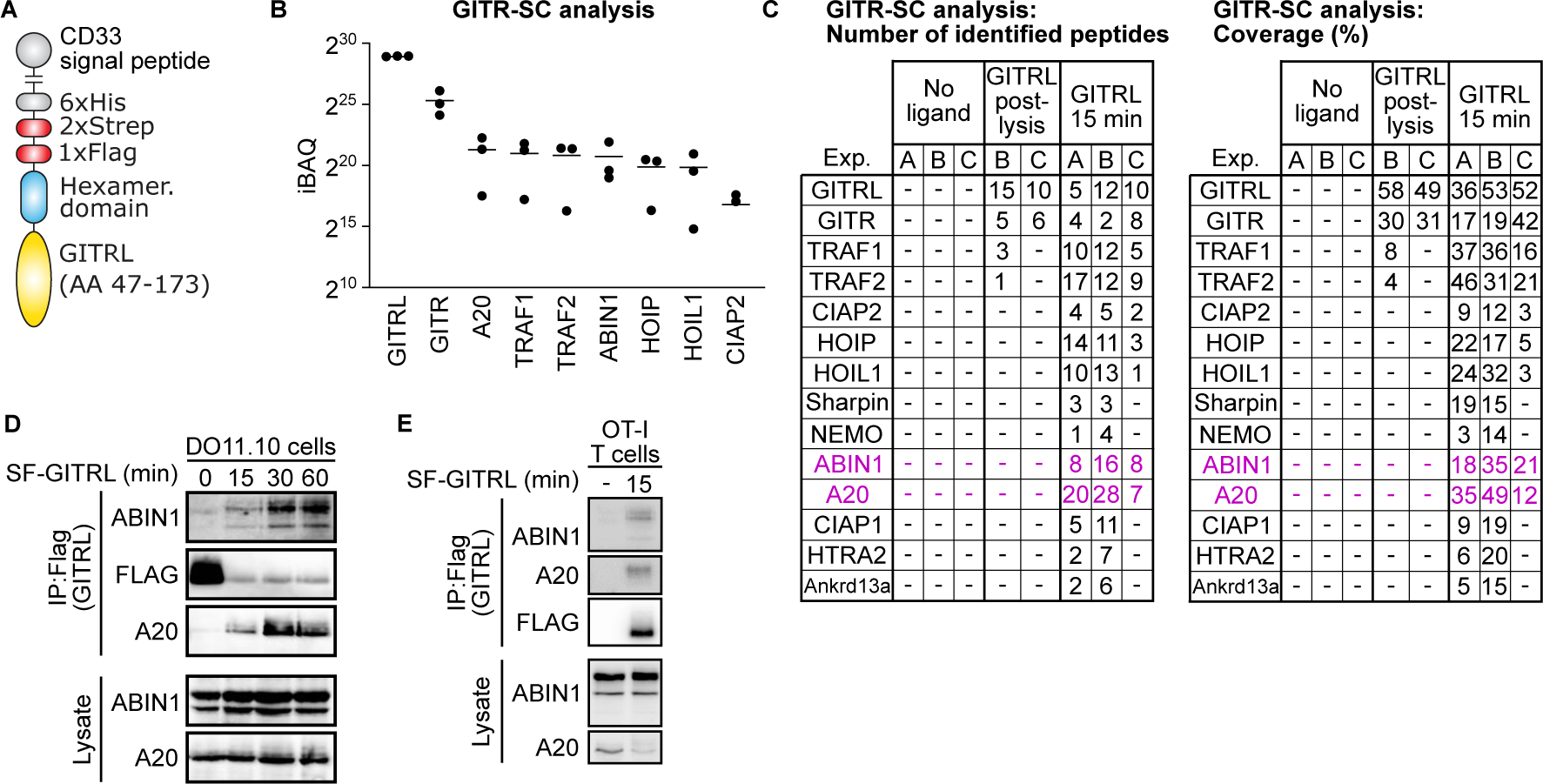
Analysis of the proximal GITR signaling complex (SC). **(A)** A schematic depiction of the recombinant GITRL for affinity purification. **(B-C)** Primary murine T cells were pre-activated with PMA/ionomycin for 72 hours and stimulated with the recombinant GITRL for 15 min. The cells were lysed and GITR-SC was isolated via tandem affinity purification and analyzed by mass spectrometry. iBAQ score of GITR-SC proteins identified in the in all three independent experiments (B) and the number of peptides and coverage of GITR-SC proteins identified in at least two independent experiments out of three in total (C) are shown. **(D-E)** DO11.10 cells (D) or OT-I T cells pre-activated with PMA/ionomycin (E) were activated with the GITRL followed by immunoprecipitation and immunoblotting using antibodies to indicated proteins. Representative experiments out of 2 in total are shown.

### Mouse models to study the role of ABIN1 in T cells

Based on the previous results, we hypothesized that ABIN1 is a negative regulator of T-cell activation. To study the role of ABIN1 in T cells, we aimed to establish a mouse model of ABIN1-deficiency. First, we used a mouse bearing a “knock-out first” gene trap (GT) *Abin1* allele (Fig. EV2A). Comparison of *Abin1*^WT/WT^ and *Abin1*^GT/GT^ showed that *Abin1*^GT/GT^ mice lack the full-length form ABIN1, but still express its low molecular weight form in T cells (Fig. EV2B). By a two-step crossing of the *Abin1*^GT/GT^ mice to CAG-Flp mice and then *Act*-CRE mice, we generated exon 5 lacking (dE5) whole-body *Abin1^dE5/dE5^* knock-out mice (Fig. EV2A). Contrary to our predictions, the *Abin1*^dE5^ allele was expressed as a truncated form of ABIN1 in T cells (Fig. EV2B). Both *Abin1*^GT/GT^ and *Abin1^dE5/dE5^* mice were weaned at a sub-mendelian ratio (Fig. EV2C), suggesting pre-weaning lethality with incomplete penetrance.

We immunophenotyped *Abin1*^GT/GT^ and *Abin1^dE5/dE5^*mice by flow cytometry (Appendix Figure 1-2). Whereas the *Abin1^dE5/dE5^*did not show any apparent phenotype in T-cell and B-cell compartments (Appendix Figure 3), we observed elevated CD4^+^ and CD8^+^ T-cell numbers and altered subset frequencies in the *Abin1^GT/GT^* mice (Appendix Figure 4). Overall, we hypothesized that the truncated ABIN1 in *Abin1^dE5/dE5^* mice preserves its function in the immune system, whereas *Abin1*^GT^ is a hypomorphic allele. To address this hypothesis, we crossed the *Abin1*^GT/GT^ mice with the *Act*-CRE mice to generate the *Abin1*^GTKO^ allele lacking exon 5 and still carrying part of the GT cassette (Fig. EV2A). *Abin1*^GTKO/GTKO^ mice lacked both natural forms of ABIN1, did not express the truncated variant typical for *Abin1*^dE5^ allele (Fig. EV2B) and were born in a sub-mendelian ratio (Fig. 2A). The *Abin1*^GTKO/GTKO^ mice showed an altered T-cell and B-cell compartments similar to *Abin1*^GT/GT^ (Fig. 2B, Fig. EV2D-E, Appendix Figure 4), indicating that the *Abin1*^GT^ and *Abin1*^GTKO^ are null or severely hypomorphic alleles and that the *Abin1*^dE5^ allele does not lead to functional ABIN1 deficiency in lymphocytes. For this reason, we decided to continue with the *Abin1*^GTKO/GTKO^ mice.

**Figure 2.**
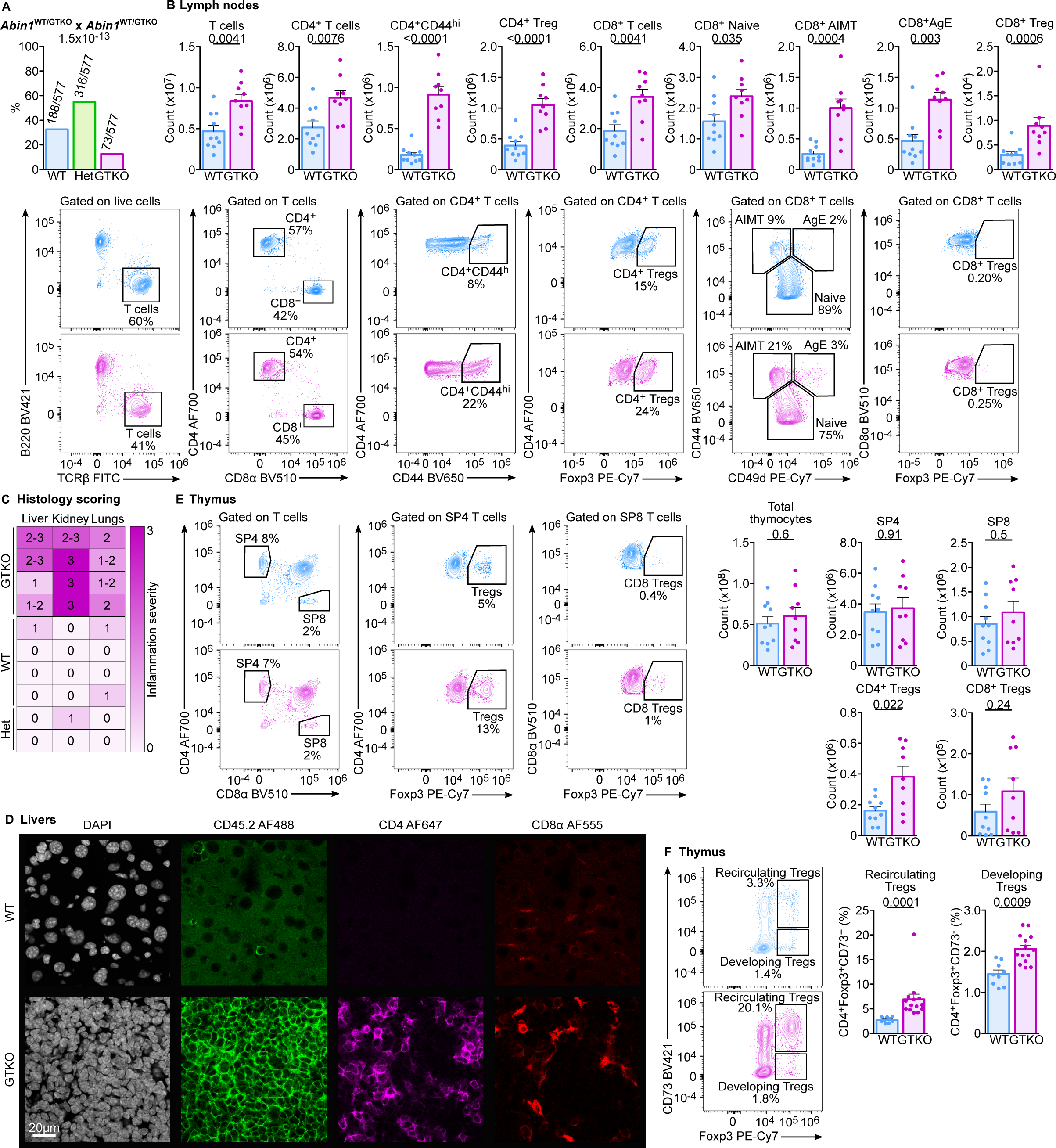
Characterization of the *Abin1*^WT/WT^ (WT) and *Abin1*^GTKO/GTKO^ (GTKO) mice. **(A)** Heterozygous *Abin1^WT/GTKO^* (HET) mice were mated and the genotype of their offspring was determined upon weaning. The frequencies and numbers of pups with particular genotypes are indicated. n = 577 offspring mice in total from 18 breedings. **(B)** Lymph node cells were stained with indicated antibodies and analyzed by flow cytometry. Representative dot plots and aggregate counts of indicated subsets are shown. n = 10 (WT) or 9 (GTKO) mice in 6 independent experiments. **(C)** Blinded histological scoring of indicated organs of 20-26 week old mice based on H&E staining. 0 – no pathology, 3 – very strong leukocyte infiltration and tissue damage. n = 4 (WT and GTKO) or 2 (HET). **(D)** Cryosections of liver of WT and GTKO mice were stained with indicated antibodies and DAPI (nuclei) and analyzed by confocal fluorescence microscopy. Representative sections out of 4 mice per group in total. **(E)** Fixed and permeabilized thymocytes from WT and GTKO mice were stained with indicated antibodies and analyzed by flow cytometry. Representative dot plots and counts of indicated subsets are shown. n = 10 (WT) or 9 (GTKO) in 6 independent experiments. **(F)** Fixed and permeabilized thymocytes from WT and GTKO mice were stained with indicated antibodies and analyzed by flow cytometry. SP4 thymocytes were gated. Representative dot plots and the frequencies of indicated subsets among SP4 cells are shown. n = 9 (WT) or 14 (GTKO) in 7 independent experiments. When applicable, the results are shown as means + SEM and p-values are indicated. Statistical significance was determined by a binomial test (A) or two tailed Mann-Whitney test (B, E, G). AIMT – antigen inexperienced memory-like T cells; AE – antigen experienced cells

**Figure 3.**
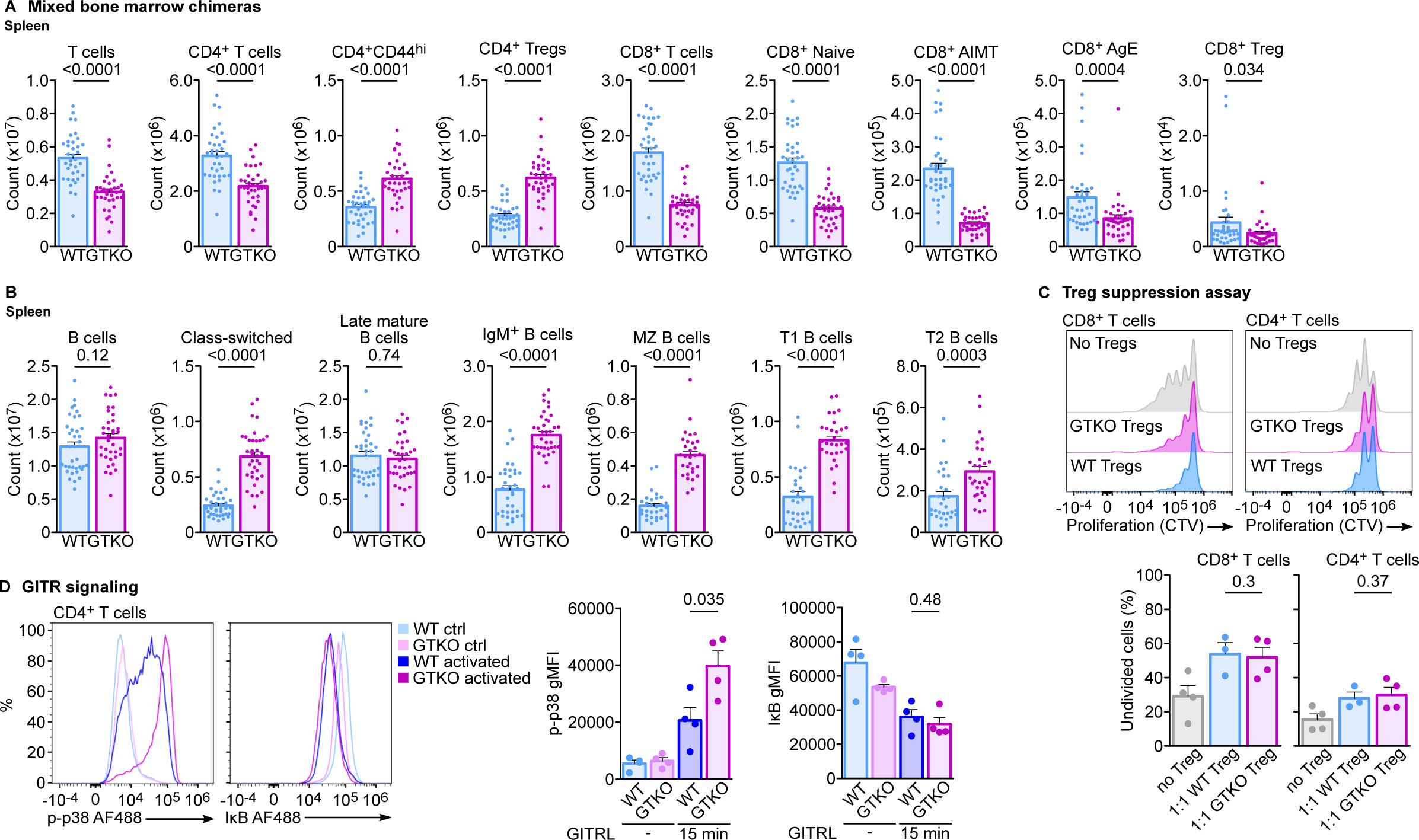
Intrinsic roles of ABIN1 in T cells. **(A, B)** Mixed bone marrow chimeras generated by transplanting Ly5.1/Ly5.2 *Abin1*^WT/WT^ (WT) and Ly5.2 *Abin1*^GTKO/GTKO^ (GTKO) bone marrow cells into irradiated Ly5.1 WT hosts, which were analyzed after 8 weeks post transplantation. Splenocytes were stained with the indicated antibodies and analyzed by flow cytometry. Counts of indicated subsets of T cells (A) and B cell subsets (B) are shown. n = 36 mice in 5 independent experiments. **(C)** CD4^+^ or CD8^+^ T cells were labeled with Cell Trace Violet dye (CTV) and FACS sorted. CD4^+^ GFP^+^ (FOXP3^+^) Treg cells were FACS sorted from DEREG^+^ *Abin1*^WT/WT^ or DEREG^+^ *Abin1*^GTKO/GTKO^ mice. CTV-loaded CD4^+^ or CD8^+^ T cells were mixed with *Abin1*^WT/WT^ Treg or *Abin1*^GTKO/GTKO^ Treg cells at 1:1 ratio. Cells were co-cultured for 72 hours and their proliferation was measured by flow cytometry. As a control, CTV-loaded T cells were cultured alone. Representative histograms and the quantification of undivided cells are shown. n = 3 (WT) or 4 (GTKO) mice in 4 independent experiments. **(D)** PMA/ionomycine pre-activated lymph node cells from WT or GTKO mice were activated with GITRL or left untreated (controls). Indicated signaling intermediates in CD4^+^ and CD8^+^ T cells were analyzed by flow cytometry. Representative histograms and aggregate results of phospho-p38 and IκB levels in CD4^+^ T cells. n = 4 independent experiments. When applicable, the results are shown as means + SEM and p-values are indicated. Statistical significance was determined by two-tailed Mann-Whitney test (A-B) or two-tailed Student’s t test (C-D). AIMT – antigen inexperienced memory-like T cells; AE – antigen experienced cells

**Figure 4.**
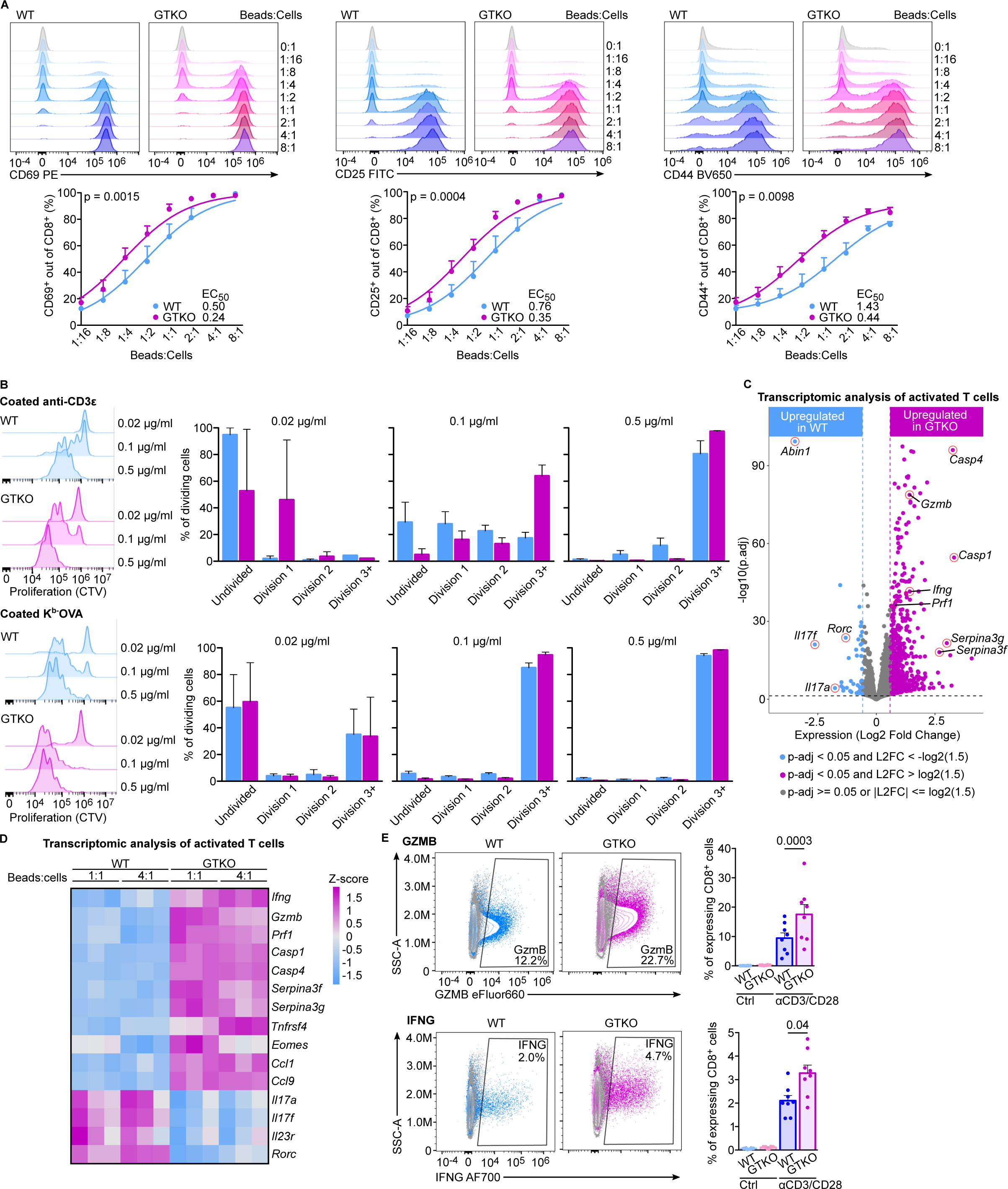
ABIN1-deficient T cells are hyperresponsive. **(A)** FACS-sorted CD8^+^ T cells from OT-I *Rag2*^KO/KO^ *Abin1*^WT/WT^ (WT) or *Abin1*^GTKO/GTKO^ (GTKO) mice were activated with anti-CD3/CD28 beads at indicated ratios for 16h. CD25, CD44, and CD69 were stained with antibodies and the percentage of positive cells was determined by flow cytometry. Representative histograms and aggregate results with log(agonist) vs. response nonlinear regression fits (three parameters) and EC_50_ are shown. n = 3 independent experiments. **(B)** Splenocytes and lymph node cells from WT and GTKO mice were FACS-sorted and labeled with Cell Trace Violet Dye. Subsequently, the cells were cultivated in plates coated with indicated concentrations of anti-CD3 antibody or monomeric K^b^-OVA for 72 h. Cell proliferation was analyzed by flow cytometry. Representative histograms and quantified frequencies in individual peaks are shown. n = 4 independent experiments. **(C-D)** Lymph node cells from WT or GTKO mice were activated with anti-CD3/CD28 beads at 1:1 or 1:4 ratios for 16 h prior to RNA isolation and RNA sequencing. n = 3 biological replicates. **(C)** A volcano plot. **(D)** A heatmap showing the differential expression of selected genes**. (E)** Lymph node cells from WT and GTKO mice were activated with anti-CD3/CD28 beads at 1:1 ratio for 16 h. Expression of GZMB and IFNG was analyzed by flow cytometry. Representative dot plots and aggregate results for indicated groups are shown. n = 8 (WT and GTKO) mice in 3 independent experiments. When applicable, results are shown as means + SEM and p-values are indicated. Statistical significance was determined using the extra sum of squares F test (A), or by two-tailed Mann-Whitney test (E).

### ABIN1-deficiency alters the lymphocyte compartment in extrinsic and intrinsic manners

*Abin1*^GTKO/GTKO^ had signs of autoimmune pathology demonstrated by tissue damage and infiltration of immune cells, mostly T cells, in the liver, kidney, and lungs (Fig. 2C-D, Fig. EV2F-G). Accordingly, *Abin1*^GTKO/GTKO^ showed altered lymphocyte compartment compared to *Abin1*^WT/WT^ (WT) mice, including expanded peripheral CD8^+^ and CD4^+^ T cells (Fig. 2B). A higher percentage of CD8^+^ T cells had phenotypes of CD44^+^ CD49d^+^ antigen-experienced (AE) or CD44^+^ CD49d^-^ antigen inexperienced memory-like T cells (AIMT) in *Abin1*^GTKO/GTKO^ than in the WT mice. Among CD4^+^ T cells, the *Abin1*^GTKO/GTKO^ mice had high frequencies of CD44^+^ activated cells and FOXP3^+^ regulatory T cells (Tregs).

The thymic development of T cells was comparable in the WT and *Abin1*^GTKO/GTKO^ mice (Fig. 2E, Fig. EV2H), indicating that the difference in the T-cell compartment occurs in mature T cells in the periphery. The only notable difference in the thymus was the abundance of CD4^+^ FOXP3^+^ Tregs. Based on their expression of CD73 (Fig. 2F) [19], we concluded that these additional Tregs recirculated to the thymus from the periphery.

To discriminate the intrinsic and extrinsic phenotypes of ABIN1-deficiency, we generated mixed bone marrow chimeras using a 1:1 mixture of Ly5.2 *Abin1*^GTKO/GTKO^ and congenic Ly5.1/Ly5.2 WT bone marrow donor cells transplanted into irradiated Ly5.1 WT hosts (Fig. 3A, Fig. EV3A, Appendix Fig. 2). We did not observe enhanced expansion and differentiation of *Abin1*^GTKO/GTKO^ T cells in these chimeras, suggesting that the steady-state changes in this compartment in the *Abin1*^GTKO/GTKO^ mice were largely extrinsic. The count of *Abin1*^GTKO/GTKO^ peripheral T cells was even slightly lower than of WT T cells in this competitive setup (Fig. 3A, Fig. EV3A). In contrast, the enhanced formation of Treg cells (Fig. 3A Fig. EV3A) and the enhanced isotype switching of B cells and their differentiation into plasma cells (Fig. 3B Fig. EV3B) were intrinsic phenotypes caused by the ABIN1 deficiency. The *Abin1*^GTKO/GTKO^ Tregs were functional, as they had a comparable suppression capacity to Tregs from the WT mouse (Fig. 3C).

### ABIN1 is a negative regulator of co-stimulation and antigenic signaling in T cells

To study the role of ABIN1 in GITR signaling, we activated T cells from the *Abin1*^GTKO/GTKO^ and WT mice with PMA/ionomycin overnight and then, the PMA/ionomycin was then washed out and the cells were cultured with IL-2 for 72h to induce the expression of GITR (Fig. EV1A). Next, we stimulated them with GITRL. The GITR signaling triggered the NF-κB pathway (degradation of IκB) and p38 MAPK pathway (phosphorylation of p38) in CD4^+^ and to lower extent in CD8^+^ T cells (Fig. 3D, Fig. EV3C). The ABIN1 deficiency leads to slightly lower basal IκB levels in the non-activated T cells, but did not alter the response to GITRL stimulation (Fig. 3D, Fig. EV3C). In contrast, p38 phosphorylation was elevated in CD4^+^ *Abin1*^GTKO/GTKO^ cells in comparison to WT cells, indicating that ABIN1 is a negative regulator of the GITR-p38 axis (Fig. 3D). We did not observe this effect in CD8^+^ T cells (Fig. EV3C). Overall, these results revealed the negative role of ABIN1 in the co-stimulation signaling, although the effect observed in this ex vivo assay was limited to CD4^+^ T cells and the p38 pathway.

To remove the extrinsic effects of ABIN1-deficiency such as the autoimmune enviroment, we crossed the *Abin1*^GTKO/GTKO^ mice to OT-I *Rag2*^KO/KO^ (henceforth OT-I) mice, which lack B cells and the only T cells formed are H-2K^b^-SIINFEKL (OVA)-specific CD8^+^ T cells. The *Abin1*^GTKO/GTKO^ and WT OT-I mice showed a comparable thymic development and formed OT-I T cells with predominant naïve CD44^-^ CD49d^-^ phenotype (Fig. EV4A). The only clear differences between the strains were higher frequencies of mature SP8 thymocytes and TCR-negative splenocytes, perhaps innate lymphocytes, in the *Abin1*^GTKO/GTKO^ OT-I mice (Fig. EV4A).

To examine the role of ABIN1 in the antigenic responses of primary T cells, we activated T cells isolated from *Abin1*^GTKO/GTKO^ and WT mice using anti-CD3/CD28 beads. The ABIN1-deficient T cells showed stronger upregulation of CD44, CD69, and CD25 than WT cells (Fig. 4A). In the next step, we activated *Abin1*^GTKO/GTKO^ and WT OT-I T cells with plate-bound anti-CD3 antibody or K^b^-OVA monomer. In both cases, *Abin1*^GTKO/GTKO^ OT-I T cells showed more rapid proliferation, especially to the suboptimal concentration of the agonist (Fig. 4B).

In the next step, we activated the *Abin1*^GTKO/GTKO^ and WT OT-I cells with anti-CD3/CD28 beads to analyze their transcription profile using RNA sequencing. ABIN1-deficient OT-I T cells triggered a different activation-induced reprogramming than WT OT-I T cells (Fig. EV4B). Activated *Abin1*^GTKO/GTKO^ OT-I T cells showed higher expression of some genes, including those encoding effector molecules (*Gzmb* and *Ifng*), caspases (*Casp1* and *Casp4)* and protease inhibitors (*Serpina3f*, *Serpina3g*) (Fig. 4C-D, Supplemental Table 1). On the other hand, ABIN1-deficient T cells expressed lower levels of *Il17a*, *Il17f*, and *Il23r*, which are genes typical for unconventional Tc17 cells (Fig. 4C-D) [20]. Accordingly, ABIN1-deficient T cells expressed higher levels of a transcription factor *Eomes*, but lower levels of *Rorc*, suggesting that ABIN1 suppresses the conventional cytotoxic effectors differentiation program, but promotes the Tc17 effectors. The elevated expression of key effector molecules GZMB and IFNG in activated *Abin1*^GTKO/GTKO^ OT-I T cells was confirmed by flow cytometry (Fig. 4E). Overall, these data suggested that ABIN1 is a negative regulator of TCR-mediated activation and expression of the key effector molecules in primary CD8^+^ T cells.

### ABIN1 regulates TCR-mediated p38 activation

To investigate the role of ABIN1 during T-cell immune responses in vivo, we observed that freshly isolated and immediately fixed steady-state *Abin1*^GTKO/GTKO^ OT-I T cells have higher levels of p38 phosphorylation than WT OT-I T cells (Fig. 5A). To assess the response of individual signaling pathways, we activated the *Abin1*^GTKO/GTKO^ and WT OT-I T cells using T2-Kb cell line loaded with low (0.1 nM) or high (1 μM) concentration of OVA peptide and measured downstream signaling intermediates by flow cytometry and flow imaging. We observed slightly elevated phosphorylation of p38, but not ERK MAPK kinase (Fig. 5B) in *Abin1*^GTKO/GTKO^ OT-I T cells. Moreover, we observed that ABIN1-deficient T cells showed slightly augmented nuclear translocation of NF-κB, but only very minor, if any, effect on NFAT nuclear translocation (Fig. 5C). Overall, these results suggested that ABIN1 selectively regulates specific downstream signaling cascades leading to T-cell activation, particularly p38 and NF-κB pathways.

**Figure 5.**
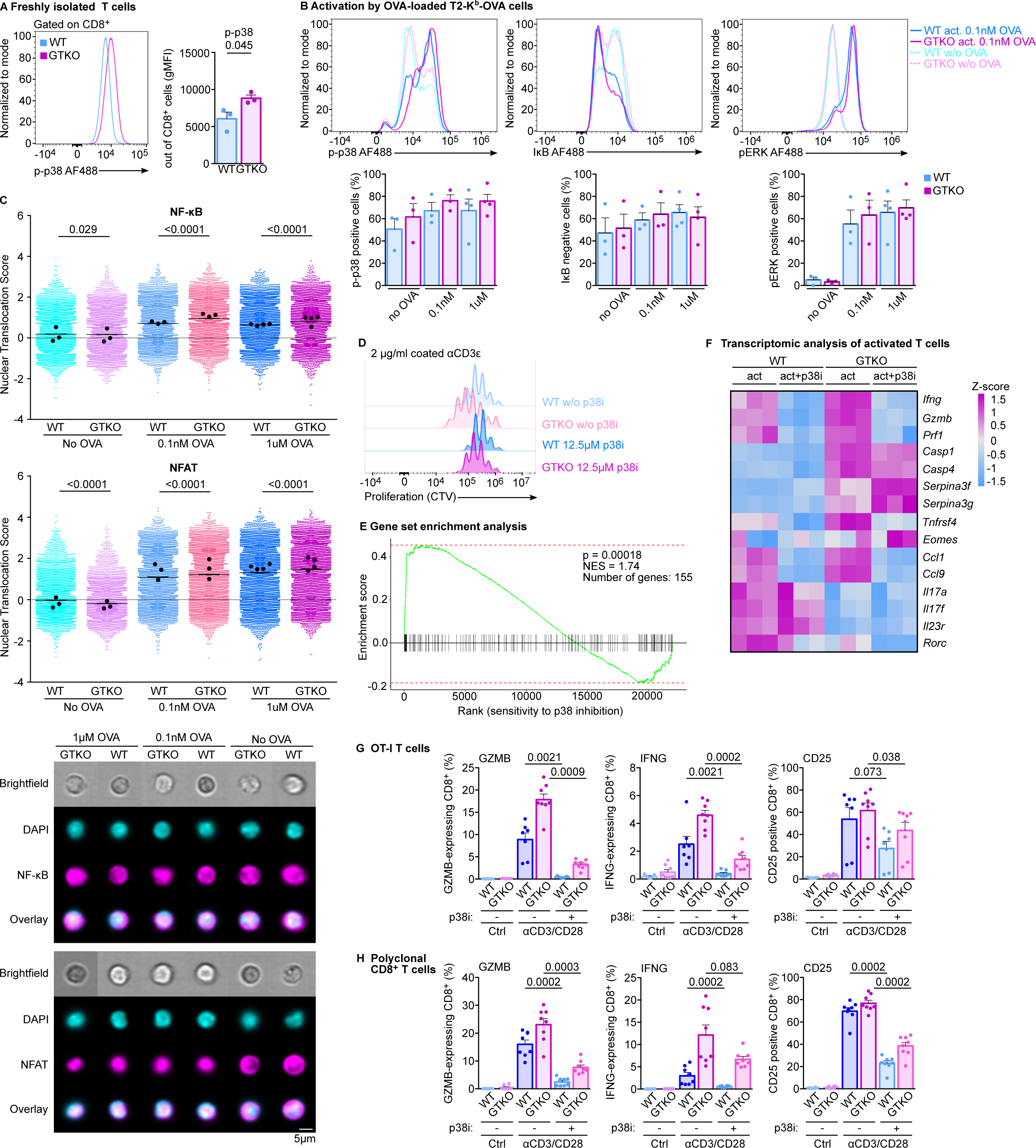
ABIN1 regulates p38 and NFκB signaling pathways. **(A)** Freshly isolated and immediately fixed and permeabilized splenocytes from *Abin1*^WT/WT^ OT-I *Rag2*^KO/KO^ (WT) and *Abin1*^GTKO/GTKO^ OT-I *Rag2*^KO/KO^ (GTKO) were stained with anti-phospho-p38 antibody and analyzed by flow cytometry. A representative histogram and quantified geometric mean fluorescence intensity in CD8^+^ T cells are shown. n = 3 independent experiments. **(B-C)** OT-I T cells were magnetically sorted (B) or FACS-sorted (C) from the WT and GTKO mice. T2-Kb cells were loaded with indicated concentrations of OVA peptide. Subsequently, CD8^+^ T cells were activated with T2-Kb cells at 2:1 ratio. (B) Activation of indicated signaling pathways was analyzed by flow cytometry. Representative histograms and aggregate results of the frequencies of positive cells are shown. n = 4 independent experiments. (C) Nuclear translocations of NF-κB and NFAT were analyzed by imaging flow cytometry. The plots shows nuclear translocations scores of pooled cells from 3 (no OVA, 0.1 nM OVA) or 4 (1 μM OVA) independent experiments (2000 cells per experiment) with indicated medians of each experiment (black dot). Representative images are shown. **(D)** Lymph node cells from WT and GTKO mice were labeled with Cell Trace Violet Dye. Subsequently, the cells were cultivated in anti-CD3 coated plates for 72 h with or without p38 MAPK inhibitor SB203580 (12.5 µM) and analyzed by flow cytometry. Representative histograms are shown. n = 4 independent experiments. **(E-F)** Lymph node cells from WT or GTKO mice were activated with anti-CD3/CD28 beads at 1:1 ratio with or without the p38 MAPK inhibitor (12.5 µM) for 16h prior to RNA isolation and RNA sequencing. n = 3 biological replicates. **(E)** A gene set enrichment analysis using a set of genes which were upregulated in GTKO vs WT T cells upon anti-CD3/CD28 in this and the previous set of experiments (Figure 4C-D, Supplemental Table 3) and genes ranked according to their sensitivity to p38 inhibition upon activation (irrespective of the genotype). **(F)** A heatmap showing the differential expression of selected genes. **(G-H)** Lymph node cells from OT-I WT and GTKO (G) or polyclonal WT and GTKO (H) mice were activated with anti-CD3/CD28 beads at 1:1 ratio with or without the p38 MAPK inhibitor (12.5 µM) for 16h. Expression of GZMB, IFNG, and CD25 was analyzed by flow cytometry. The frequencies of positive cells are shown. (G) n = 7 (WT), or 8 (GTKO) mice in 3 independent experiments. (H) n = 8 mice in 3 independent experiments. When applicable, results are shown as means + SEM and p-values are indicated. Statistical significance was determined using unpaired t-test (A), or two-tailed Mann-Whitney test (B, C, G).

To address whether the differential activation of p38 in *Abin1*^GTKO/GTKO^ and WT OT-I T cells could explain the differences in their ex vivo response to the antigen (Fig. 4), we activated T cells by immobilized anti-CD3 antibody in the presence or absence of p38 inhibitor. The proliferation advantage of *Abin1*^GTKO/GTKO^ T cells was largely diminished by p38 inhibition (Fig. 5D). In the next step, we activated *Abin1*^GTKO/GTKO^ and WT OT-I T cells with anti-CD3/CD28 beads in the presence or absence of the p38 inhibitor and performed the transcriptomic analysis by RNA sequencing (Fig. EV5A, Supplemental Table 2). The analysis of samples without the p38 inhibitor corresponded well to the results of the previous experiment (Fig. EV5B-C). We generated a set of genes which were upregulated in *Abin1*^GTKO/GTKO^ T cells in both experiments (Supplemental Table 3). Using a gene set enrichment analysis, we observed that many genes upregulated in activated *Abin1*^GTKO/GTKO^ T cells are sensitive to the inhibition of the p38 pathway (Fig. 5E). The genes upregulated in ABIN1-deficient T cells and down-regulated upon p38 inhibition in activated *Abin1*^GTKO/GTKO^ vs. WT cells included *Gzmb*, *Ifng*, and *Tnfrsf4* (OX40), but not *Casp1* and *Casp4* (Fig. 5F, Fig. EV5D). The down-regulation of Tc17-associated genes, was not sensitive to p38 inhibition (Fig. 5F). To verify the transcriptomic data on a protein level, we detected GZMB and IFNG in *Abin1*^GTKO/GTKO^ and WT OT-I or polyclonal T cells activated by αCD3/CD28 beads in the presence or absence of p38 inhibitor. We observed that p38 inhibition decreases the levels of GZMB and IFNG in monoclonal and polyclonal CD8^+^ T cells (Fig. 5G-H, Fig. EV5E-F). Moreover, the activation-induced expression of CD25 was also sensitive to p38 inhibition (Fig. 5G-H, Fig. EV5E-F).

Overall, these data reveal ABIN1 as a negative regulator of p38 activation in T cells, which leads to upregulation of p38-dependent key effector genes in ABIN1-deficient T cells.

### ABIN1 is a negative regulator of cytotoxic T-cell responses in vivo

We adoptively transferred *Abin1*^GTKO/GTKO^ or WT OT-I T cells into congenic Ly5.1 recipients, which were subsequently infected with *Listeria monocytogenes* expressing OVA (Lm-OVA) or its lower affinity variant Q4H7 (Lm-Q4H7). We observed higher expansion, preferential differentiation of *Abin1*^GTKO/GTKO^ OT-I T cells into KLRG1^+^ IL7R^-^ short-lived effector T cells (SLEC) at the expense of KLRG1^-^ IL7R^+^ memory precursors and higher expression of CD25 in comparison to the WT controls upon Lm-OVA infection (Fig. 6A, Fig. EV6A, Appendix Figure 2). The hyper-responsiveness of *Abin1*^GTKO/GTKO^ T cells was pronounced in the infection with *Listeria* expressing an altered suboptimal OT-I antigen Q4H7 (Lm-Q4H7) (Fig. 6B, Fig. EV6B). We obtained very similar results with *Abin1*^GT/GT^ OT-I T cells (Fig. EV6C), i.e., T cells bearing the original GT version of the ABIN1 targeted allele (Fig. EV2A). The enhanced formation of SLEC cells in *Abin1*^GTKO/GTKO^ OT-I T cells was also observed upon the infection with LCMV-OVA (Fig. EV6D).

**Figure 6.**
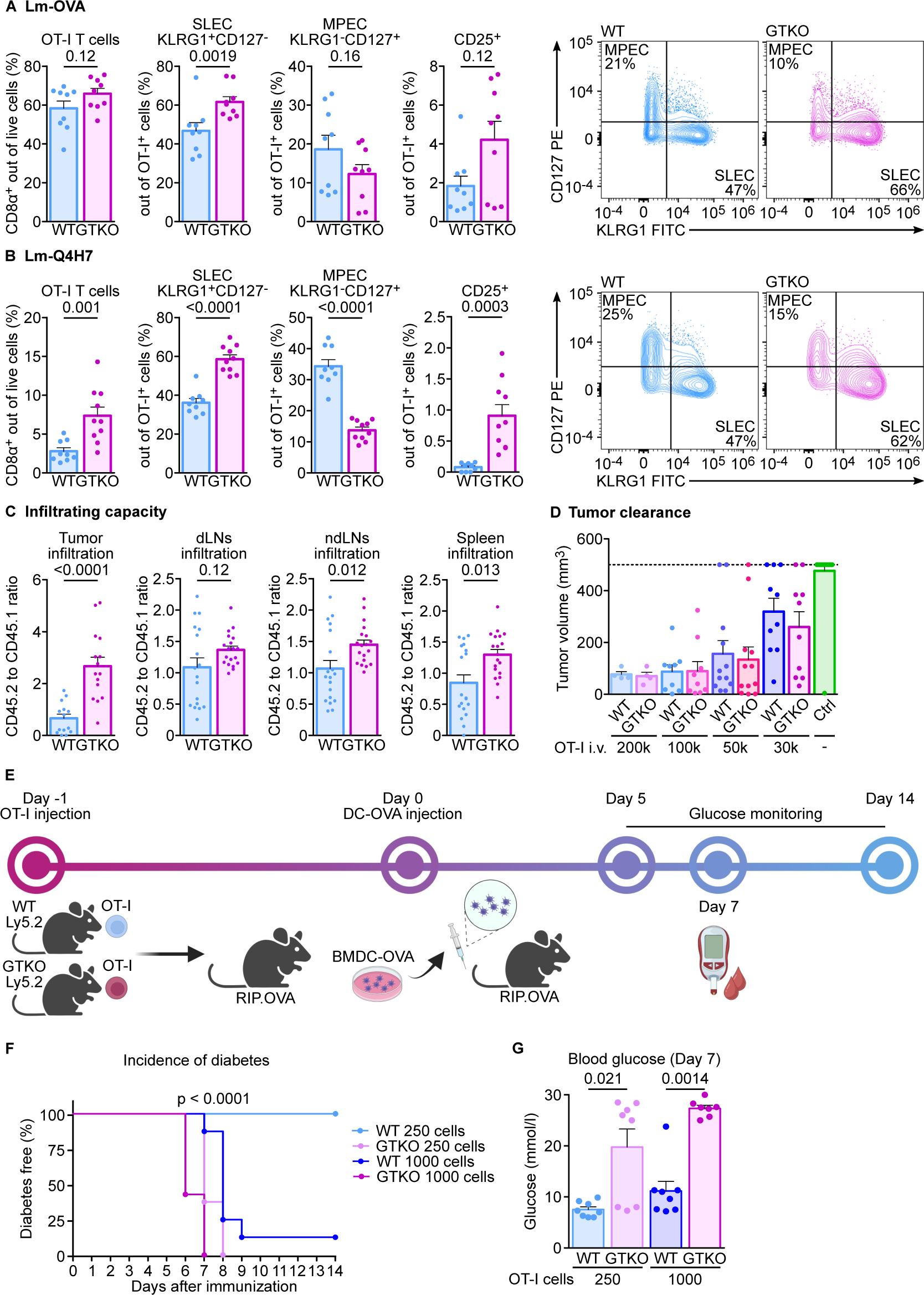
ABIN1 regulates T-cell responses in vivo. **(A-B)** 1×10^5^ T cells from *Abin1*^WT/WT^ OT-I *Rag2*^KO/KO^ (WT) or *Abin1*^GTKO/GTKO^ OT-I *Rag2*^KO/KO^ (GTKO) were adoptively transferred to Ly5.1 hosts that were infected with *Listeria monocytogenes* expressing (A) ovalbumin (Lm-OVA) or (B) its lower-affinity variant (Q4H7). Splenocytes were analyzed by flow cytometry on day 6 post-infection. Representative dot plots and frequencies of indicated T cell subsets are shown. (A) n = 9 mice per group in 3 independent experiments. (B) n = 9 (WT), or 10 (GTKO) mice in 3 independent experiments. **(C)** Lymph node cells from Ly5.1 WT OT-I *Rag2*^KO/KO^ mice were mixed at 1:1 ratio with cells from Ly5.2 WT or GTKO littermates and mixed donor cells were adoptively transferred to congenic Ly5.1/Ly5.2 WT mice bearing MC-38 tumors. On day 7 post transfer, the tumors, draining lymph nodes, non-draining lymph nodes, and spleens were analyzed by flow cytometry and the ratio of Ly5.2 (WT or GTKO) to Ly5.1 (WT) donor cells was calculated. n = 18 mice per group in 3 independent experiments. **(D)** Indicated numbers of lymph node cells from WT and GTKO (OT-I) were adoptively transferred into CD3ε^-/-^ host bearing MC-38 tumor. No cells were transferred in control group. The tumor volume was monitored for consecutive 23 days, or up to the endpoint volume of 500 mm^3^. The maximal tumor volume for each mouse is shown. n = 4 for 2×10^5^ transferred cells, 9 for 1×10^5^ transferred cells, 12 (WT and GTKO) for 5×10^4^ transferred cells, 10 for 3×10^4^ transferred cells, and 21 for no transfer in 6 independent experiments in total. **(E-G)** Indicated numbers of WT or GTKO (OT-I) cells were adoptively transferred to RIP.OVA host mice. On the following day, mice were immunized with OVA-loaded bone marrow-derived dendritic cells. The concentration of glucose in urine was monitored on a daily basis for 14 days and on day 7 glucose in blood was measured. The mouse was considered diabetic when the concentration of glucose reached 1000 mg/dl in urine. n = 8 mice per group in 2 independent experiments. (E) The scheme of the experiment. (F) Blood glucose concentration on day 7**. (G)** Survival (diabetes-free curve). When applicable, results are shown as means + SEM and p-values are indicated. Statistical significance was determined using two-tailed Mann-Whitney test (A, B, C, F) or Log-rank (Mantel-Cox) test (G).

We tested the response of *Abin1*^GTKO/GTKO^ and WT OT-I T cells to tumors, using mice with implanted subcutaneous MC38-OVA tumors. We adoptively transferred Ly5.2 *Abin1*^GTKO/GTKO^ OT-I T cells or WT OT-I T cells mixed with competitor Ly5.1 WT OT-I T cells at 1:1 ratio into MC38-OVA tumor bearing Ly5.1/Ly5.2 heterozygous mice. After seven days, we determined the relative abundance of WT and GTKO OT-I T cells in the lymphoid organs and in the tumor as a ratio of Ly5.2^+^ to Ly5.1^+^ OT-I T cells (Fig. 6C, Fig. EV6E). Whereas the ratio of WT Ly5.2 OT-I to WT Ly5.1 OT-I cells was close to one, as expected, we observed a competitive advantage of *Abin1*^GTKO/GTKO^ over the Ly5.1 WT OT-I T cells, which was most pronounced in the tumor (Fig. 6C). The strong ability of *Abin1*^GTKO/GTKO^ OT-I T cells to infiltrate the tumor motivated us to assess the anti-tumor response of these cells. We observed that both *Abin1*^GTKO/GTKO^ OT-I T cells and WT OT-I T are able to suppress the growth of the MC38-OVA tumor when transferred at high numbers. However, we did not observe that *Abin1*^GTKO/GTKO^ OT-I T cells have significantly better anti-tumor properties than their WT OT-I counterparts (Fig. 6D, Fig. EV6F).

The above-presented data indicated that ABIN1 is a negative regulator of T-cell responses in vivo and thus, might be important for maintaining T-cell peripheral tolerance. We compared the potency of *Abin1*^GTKO/GTKO^ and WT OT-I T cells to induce experimental autoimmune diabetes model. The model is based on a transfer of OT-I T cells into a transgenic RIP.OVA mouse expressing OVA under the rat insulin promoter [21] followed by their in vivo priming with bone marrow-derived dendritic cells loaded with OVA [22]. We observed that ABIN1-deficient OT-I T cells are much more potent in inducing this autoimmune pathology (Fig. 6E-G), suggesting that *Abin1*^GTKO/GTKO^ T cells have a supraphysiological ability to undergo antigen-induced expansion, effector cell formation, tissue infiltration, and target cell killing, which releases them from the Treg-mediated control [22] in this autoimmune model.

Altogether, our data identify ABIN1 as an intrinsic negative regulator of CD8^+^ T-cell responses via inhibiting antigen-induced and co-stimulatory signaling cascades, especially the p38 MAPK pathway.

## Discussion

We have identified ABIN1 as a part of the SCs of two T-cell co-stimulation receptors, GITR and OX40. Most likely, ABIN1 is recruited to the complex via its binding to linear polyubiquitin chains generated by the LUBAC ubiquitin ligase [23] also found in the SCs of GITR. LUBAC is a signal transducer in TNFR1 and MYD88 signaling [24]. Recently, LUBAC subunits were identified in the SC of CD137 TNFR superfamily receptor [6], but its involvement in GITR signaling was not shown before. We speculate that the recruitment of LUBAC, which generates linear polyubiquitin chains linear polyubiquitin chains and activates NEMO to trigger downstream NF-κB activation, is a common mechanism of signaling of T-cell co-stimulation receptors from the TNFR superfamily [25]. The recent report analyzed the SC of CD137 to find components largely overlapping with our detection of the GITR SC (LUBAC, cIAP2, TRAF1, TRAF2, A20) [6]. Although the LUBAC complex was present in the SC of CD137, the authors did not detect ABIN1 [6], presumably because of technical reasons. Recently, ABIN1 was reported to interact with the CARD11-CBM complex upon TCR signaling and to inhibit the TCR-induced NFκB activation in a human T-cell line [15], indicating that ABIN1 regulates antigenic as well as co-stimulation signaling in T cells.

Since ABIN1 is involved in multiple T-cell signaling pathways and its role in T cells in vivo has not been addressed, we characterized T cells in ABIN1-deficient mice. Our ABIN1^GTKO/GTKO^ mouse model largely recapitulated previous models of ABIN1-deficiency, such as partial pre-weaning lethality, intrinsic hyperactivation of B cells, and tissue pathology [10, 11]. We observed hyperactivation of T cells and their infiltration into tissues in these mice. The experiments with bone marrow chimeras showed that these T-cell phenotypes are driven by extrinsic factors, plausibly by enhanced TNF-mediated cell death and subsequent tissue damage. On the contrary, the enhanced formation of regulatory CD4^+^ T cells in *Abin1*^GTKO/GTKO^ mice, also previously shown for ABIN1 knock-in mice with disrupted ubiquitin binding [11], is a T-cell intrinsic phenotype. We crossed the *Abin1*^GTKO/GTKO^ mice to monoclonal OT-I *Rag2*^KO/KO^ mice, completely lacking the B-cell compartment and T-cell diversity, to limit the T-cell extrinsic phenotype. Using these mice, we compared WT and ABIN1-deficient OT-I T cells in ex vivo assays and in experiments based on the adoptive transfer into WT hosts.

*Abin1*^GTKO/GTKO^ OT-I T cells exhibited enhanced responses to antigenic and anti-CD3/anti-CD28 induced response ex vivo. Moreover, they showed upregulation of key effector molecules GZMB and IFNG. We observed slightly upregulated NFκB and p38 pathways and virtually undetectable effects on the ERK and NFAT pathways upon the ex vivo activation. The inhibition of p38 resulted in down-regulation of a significant proportion of genes, which were also up-regulated in the *Abin1*^GTKO/GTKO^ OT-I T cells upon activation, including GZMB and IFNG.

To our knowledge, the fact that the p38 pathways up-regulates GZMB upon TCR signaling was not previously shown, although there is a report that p38 controls GZMB expression in neutrophils in rats [26]. The upregulation of IFNG by p38 was previously demonstrated for CD8^+^ as well as CD4^+^ T cells [27, 28]. It was also proposed that the p38 pathway contributes to the T-cell apoptosis [27] and senescence [29, 30]. Although we did not specifically focus on this question, the ∼2 fold lower numbers of *Abin1*^GTKO/GTKO^ than WT CD8^+^ T cells in periphery of the mixed bone marrow chimeras suggest, that the enhanced apoptosis might be the cause. A recent study shows that p38 inhibition in cultured activated murine T cells leads to upregulation of memory-signature genes (*Tcf7*) and down-regulation of effector genes (*Klrg1*, *Gzmc, Prf1, Havcr2*) [31], which is essentially in line with our in vitro and in vivo data with *Abin1*^GTKO/GTKO^ and p38-inhibited T cells. Although not mentioned in that article, their data show down-regulation of *Ifng* and *Gzmb* upon p38 inhibition in these cells, which corresponds with our data [31]. However, they also showed that deletion of p38 *Mapk14* improves the anti-cancer properties in adoptive T-cell therapy in mice and enhances proliferation and expression of effector molecules in human T cells [31]. In a striking contrast to theirs and ours murine data, this study also describes enhanced IFNG production upon p38 inhibition in human CD8^+^ T cells [31], indicating a species-dependent or context-dependent role of p38. Overall, these discrepancies uncover that the role of p38 in T-cell biology is incompletely understood.

p38 is activated via TAB2/3-TAK1 complex, which is assembled at K63 polyubiquitin chains in multiple signaling pathways [32]. These K63 polyubiquitin chains are formed by TRAF6 ubiquitin ligase in the CMB complex and plausibly by TRAF2/cIAP2 in the GITR complex. Since ABIN1 recruits and/or activates A20 in various signaling complexes, we propose that ABIN1 might down-regulate the p38 pathway via A20, which cleaves the K63 polyubiquitin chains [33], leading to the release and deactivation of the TAB2/3-TAK1 complex. Additionally, the TCR signaling was shown to activate p38 via an alternative pathway, triggered by the direct phosphorylation of p38 by a proximal ZAP70 kinase [34]. It is unclear, whether and how this pathway might be related to the ABIN1-mediated regulation of p38. Although the p38 pathway is up-regulated in the *Abin1*^GTKO/GTKO^ T cells and seems to be at least partially responsible for the observed phenotype, other downstream signaling pathways, such as NF-κB pathway, probably also play an important role in the hyperactivation of *Abin1*^GTKO/GTKO^ T cells observed in vitro and in vivo.

We have seen that the deficiency of ABIN1 was coupled with the upregulation of key effector molecules GZMB and IFNG typical for type 1 immunity, but also with the down-regulation of *Il17a, Il17f, Il23, and Rorc*, which are typical for type 3 immunity and expressed in a subset of Tc17 cells [35]. This suggests that ABIN1 not only generally regulates T-cell activation, but it also differentially regulates the outcomes of individual signaling pathways underlying T-cell differentiation program. It should be a matter of future investigation, whether ABIN1 also regulates the differentiation of CD4^+^ helper T cells, which can form several types of effector T cells (Th1, Th2, Th17, Tfh, Treg).

Although we did not observe a superior anti-tumor activity of ABIN1-deficient T cells, it is possible that, in another context with stronger antigenic and/or co-stimulation signaling, ABIN1-deficiency would increase anti-tumor potency. This is indicated by the experiments with autoimmune diabetes, in which *Abin1*^GTKO/GKTO^ T cells were very powerful in the disease induction, probably as a consequence of their strong response to the priming by cognate inflammatory bone marrow-derived dendritic cells. One potential direction for future research is the role of ABIN1 in adoptive cancer therapies using T cells with chimeric antigen receptors consisting of a canonical TCR signaling unit (ZETA chain) and TNFR super-family signaling domain (CD137) [36], both potentially regulated by ABIN1.

Overall, we show that ABIN1 is an important regulator of CD8^+^ T-cell activation, which limits the proliferation and formation of effector T cells and maintains peripheral tolerance. ABIN1 inhibits p38 and NFκB pathways, probably via delivering A20 deubiquitinase to NEMO [15, 37] and/or by physically blocking the polyubiquitin binding sites that recruit key signaling proteins. Our findings that ABIN1 acts as a negative regulator of CD8^+^ T-cell immune response in vivo open potential research directions to study ABIN1 as a target for T-cell therapy.

## Acknowledgment

This project was supported by the Czech Science Foundation (22-18046S to OS), project National Institute of Virology and Bacteriology (Programme EXCELES, LX22NPO5103 to OS) - funded by the European Union - Next Generation EU, Charles University Grant Agency (984120 to SJ, 274323 to VU), European Union’s Horizon 2020 research and innovation programme under grant agreement No. 802878 (ERC Starting Grant FunDiT to OS), Czech Ministry of Education, Youth and Sports and the European Regional Development Fund (OP RDI CZ.1.05/2.1.00/19.0395 and LM2023036 for the Czech Centre for Phenogenomics, OP RDI BIOCEV CZ.1.05/1.1.00/02.0109), and core funding provided by the Institute of Molecular Genetics of the Czech Academy of Sciences (RVO 68378050).

We acknowledge Ladislav Cupak for technical assistance, Dagmar Zudova (Czech Centre for Phenogenomics) for the histological analysis, Karel Harant (Proteomic facility, Charles University in Prague) for proteomic analysis, Zdenek Cimburek and Matyas Sima (Flow cytometry facility, IMG) for cell sorting, and the Light Microscopy Core Facility, Institute of Molecular Genetics of the Czech Academy of Sciences, supported by Czech Ministry of Education, Youth and Sports (LM2023050 and RVO – 68378050-KAV-NPUI), for their help.

DP and ES are students of the Faculty of Science, Charles University in Prague.

## Methods

### Cell lines

DO11.10 cell line (provided by Tomas Brdicka, IMG) and T2–Kb (provided by E. Palmer, University Hospital Basel) were cultivated in RPMI media (Sigma). HEK293FT (provided by T. Brdicka, IMG) and MC-38 cells (provided by E. Palmer, University Hospital Basel) were cultivated in DMEM media (Sigma). The media were supplemented with 10% fetal bovine serum (FBS, Gibco #10270-106), 100 U/ml penicillin (BB Pharma #15/156/69-A/C), 100 µg/ml streptomycin (Sigma #59137), and 40 µg/ml gentamicin (Sandoz) and kept in 37°C, 5% CO_2_ incubator and cells were cultured in 37°C, 5% CO_2_ incubator. Cell lines were regularly tested for mycoplasma by PCR.

### Antibodies, dyes and inhibitors

Following antibodies were used for flow cytometry: anti-CD1d PE-Cy7 (clone 1B1, Biolegend #123524), anti-CD4 AF 700 (clone RM4-5, #100536), anti-CD8α BV 421 (clone 53-6.7, #100753), anti-CD8α BV 510 (clone 53-6.7, #100752), anti-CD8α FITC (clone 53-6.7, #100706), anti-CD8α PE-Cy7 (clone 53-6.7, #100722), anti-CD19 PE (clone 6D5, #115508), anti-CD23 APC (clone b3b4, #101620), anti-CD24 FITC (clone M1/69, #101806), anti-CD25 BV 605 (clone PC61, #102036), anti-CD25 FITC (clone PC61, #102006), anti-CD38 AF 488 (clone 90, #102714), anti-CD44 BV 650 (clone IM7, #103049), anti-CD45 AF 700 (clone 30-F11, #103127), anti-CD45.1 AF 700 (clone A20, #110724), anti-CD45.1 FITC (clone A20, #110706), anti-CD45.1 PerCP-Cy5.5 (clone A20, #110728), anti-CD45.2 AF 700 (clone 104, #109822), anti-CD45.2 APC (clone 104, #109814), anti-CD45.2 APC-Cy7 (clone 104, #109824), anti-CD45.2 FITC (clone 104, #109806), anti-CD45.2 PE (clone 104, #109808), anti-CD45R/B220 BV 421 (clone RA3-6B2, #103240), anti-CD49d PE-Cy7 (clone R1-2, #103618), anti-CD69 PE (clone H1.2F3, #104508), anti-CD73 BV 421 (clone TY/11.8, #127217), anti-CD127 PE (clone SB/199, #121112), anti-CD138 BV 421 (clone 281-2, #142523), anti-CD138 BV 510 (clone 281-2, #142521), anti-IgD AF 700 (clone 11-26c.2a, #405730), anti-IgM BV 421 (clone RMM-1, #406518), anti-IgM PE-Cy7 (clone RMM-1, #406513), anti-IFNγ AF 700 (clone XMG1.2, #505824), anti-KLRG1 BV 510 (clone 2F1/KLRG1, #138421), anti-KLRG1 FITC (clone 2F1/KLRG1, #138410), anti-TCRβ APC (clone H57-597, #109212), anti-TCRβ BV 711 (clone H57-597, #109243), anti-TCRβ FITC (clone H57-597, #109206), anti-FLAG APC (clone L5, #637307), FOXP3 PE-Cy7 (clone FJK-16s, eBioscience #25-5773-82), anti-GzmB eFluor660 (clone NGZB, #50-8898-82), anti-NF-κB p65 (clone C-20, Santa Cruz #sc-372), anti-IκB AF 488 (clone L35A5, Cell Signaling #5743S), anti-NFAT XP (clone D43B1, Cell Signaling #5861S), anti-pERK XP (clone D13.14.4E, Cell Signaling #4370), anti-p-p38 (clone D3F9, Cell signaling #4511). Following dyes were used for flow cytometry: Cell Trace Violet (CTV, Invitrogen #34557), Hoechst 33258 (Invitrogen #H3569), LIVE/DEAD near-IR (ThermoFisher #L34976).

Following primary antibodies were used for immunofluorescence microscopy: anti-CD4 AF 647 (clone RM4-5, Biolegend #100530), anti-CD8α (clone EPR21769, Abcam #ab217344), anti-CD45.2 AF 488 (clone 104, Biolegend #109816), DAPI (ThermoFisher #D1306).

Following primary antibodies were used for immunoblotting, cell activation: anti-CD3ε (clone 145-2C11, Biolegend #100302), anti-ABIN1 (Mrcppureagents #S345C), anti-A20 (clone A-12, Santa Cruz #sc-166692), anti-β-Actin (clone AC-15, Sigma #A1978), anti-p-p38 (clone D3F9, Cell signaling #4511), anti-DYKDDDDK Tag (clone D6W5B, Cell Signaling #14793).

Following secondary antibodies were used: goat anti-rabbit conjugated with AF 488 (Invitrogen #A-11008), AF 555 (Invitrogen #A-32732), or AF 647 (Invitrogen #A-21245), donkey anti-sheep-HRP (Jackson #713-035-147), goat anti-mouse-HRP (Jackson #115-035-146), goat anti-rabbit-HRP (Jackson #111-035-144)

P38 MAPK inhibitor SB203580 (#ab120162) was purchased from Abcam.

### Mice

All the mice had C57Bl/6J background. CD3ε^KO/KO^ (RRID:IMSR_JAX:004177) [38], DEREG (RRID: MMRRC_032050-JAX) [39], Ly5.1 (RRID: IMSR_JAX:002014) [40], OT-I Rag2^KO/KO^ (RRID:MGI:3783776, MGI:2174910) [41, 42], RIP.OVA (RRID:IMSR_JAX:005433) [21] strains were described previously.

We generated *Abin1*^GT/GT^ mice via *in vitro* fertilization of C57BL/6J oocytes with sperm carrying the GT allele (Tnip1tm1a(EUCOMM)Hmgu; RRID:IMSR_EUMMCR:1521) obtained from Infrafrontiers.

The *Abin1*^GT/GT^ mice were crossed to Cre-deleter mice Gt(ROSA)26Sor^tm1(ACTB–cre,–EGFP)Ics^ (MGI: 5285392) (*Act*-CRE) to generate *Abin1*^GTKO/GTKO^ mice missing exon 5 in *Abin1* and carrying a LacZ sequence after the exon 4 of *Abin1*. The *Abin1*^GT/GT^ mice were crossed to FLP-deleter mice Gt(ROSA)26Sor^tm2(CAG–flpo,–EYFP)Ics^ (MGI: 5285396) (CAG-Flp) to generate *Abin1*^flox/flox^ mice, followed by crossing to *Act*-CRE to cut out exon 5 to obtain *Abin1*^dE5/dE5^ mice.

Mice were bred at the Institute of Molecular Genetics of the Czech Academy of Science in a specific-pathogen-free facility. In the facility, 12h light-dark cycle was maintained, food and water ad libitum. Animal protocols were in accordance with the laws of the Czech Republic and approved by the Czech Academy of Science (identification no. 72/2017, 81/2018, 2378/2022 SOV II).

For the experiment, males and females were used. Mice were used in age 6-12 week at the beginning of the experiments. For the developmental studies, mice at age 2-3 weeks old were used. When possible, age- and sex-matched animals were used, if possible littermates were used and equally divided into experimental groups. Mice were divided into groups by their ID without prior contact with experimenter. The histology examination and one out of two diabetic experiments were blinded. Other experiments were not blinded as no subjective scoring was used.

### Preparation of knockout cell lines

To generate knock-out cell line using CRISPR/Cas9 method, gene-targeted single-guided RNA (sgRNA) was designed using chopchop software [43]. sgRNA was then inserted into pSpCas9(BB)-2A-GFP (PX458) vector [44] that was provided by Feng Zhang (Addgene plasmid #48138). List of sgRNA target sequences to prepare the ABIN1 knock-out, PAM motif in bold:

Mouse ABIN1 KO_1: 5’ – GATCCAGCGGCTCAATAAGG**TGG**
Mouse ABIN1 KO_2: 5’ – GGTGAATTCTGCTCCTCAGT**AGG**

DO11.10 cells were transfected with PX458 vector containing the sgRNA using Lipofectamine 2000 (Invitrogen #52887) according to manufacturer’s instructions. GFP^+^ cells were sorted as single cells into 96-well plate using BD Influx sorter (BD Bioscience). Obtained clones were tested via immunoblotting for expression of ABIN1 and confirmed by sequencing.

### DO11.10 cell line stimulation

Cells were kept in serum-free media for 30 min prior activation. Following, cells were stimulated with GITRL for indicated times. Subsequently, cells were lysed in 1% *n*-Dodecyl-β-D-Maltoside (DDM, Thermo Scientific #89903) in lysis buffer (30 mM Tris pH 7.4, 120 mM NaCl, 2mM KCl, 2 mM EDTA, 10% glycerol, 10 mM chloroacetamide (Sigma #C0267), 10 mM cOmplete protease inhibitor cocktail (Roche #5056489001), and PhosSTOP tablets (Roche #4906837001)). Lysates were incubated at 4°C for 30 min and cleared by centrifugation for 30 min, 2°C, 21,130 g. Samples were mixed with SDS sample buffer, reduced by 50 mM DTT, heated for 3 min at 92°C, and analyzed by immunoblotting.

### Magnetic sorting

CD4^+^ or CD8^+^ T cells were negatively sorted using biotinylated anti-B220 (clone RA3-6B2, Biolegend #103204), CD11b (clone M1/70, Biolegend #101204), and anti-CD8α (clone 2.43, ATCC# TIB-210) or anti-CD4 (clone 53-6.7, Biolegend #100704) antibodies, respectively. Subsequently, the cells were labeled with anti-Biotin MicroBeads (Miltenyi Biotec #130-090-485) and enriched using an AutoMACS Pro (Milteniy Biotech).

### Flow cytometry and cell sorting

Cells were stained with indicated antibodies for cell sorting or flow cytometry. To distinguish live cells, samples were stained with LIVE/DEAD near-IR dye (ThermoFisher #L34976) or Hoechst 33258 (Invitrogen #H3569). For staining of FOXP3 transcription factor, cells were fixed using Foxp3/Transcription Factor Staining Buffer Set (Invitrogen #00-5523-00) according to the manufacturer’s instructions. For staining of Granzyme B or IFNγ, BD Cytofix/Cytoperm Fixation/Permeabilization Kit was used (BD Bioscience #554714) according to manufacturer’s instructions. The samples were analyzed using an Aurora (Cytek), or FACSymphony (BD Bioscience). Cell sorting was done using a BD Influx or a FACSAria Ilu sorters (both BD Bioscience). FlowJo software (BD Bioscience) for used for the data analysis.

### Histology and immunofluorescence

Standard descriptive histopatology H&E staining was performed in the Czech Centre of Phenogenomics according to internal standard operating procedures. Briefly, tissue processing: automatic vacuum tissue processor (Leica ASP6025), embedding: Leica EG1150 H+C embedding station, sectioning: Leica RM2255 rotary microtome (2 µm sections), staining: Leica ST5020 + CV5030 stainer and coveslipper.

Immunofluorescent staining was done on 5 µm thick cryosections of kidney, livers, and lungs from ABIN1^WT/WT^ and ABIN1^GTKO/GTKO^ mice prepared using a cryostat (Leica CM1950). The sections on microscopy slides were fixed with 4% formaldehyde for 10 min and permeabilized with 0.1% Triton X-100 in PBS for 10 min. The samples were then blocked with 5% goat serum in PBS. Next, samples were stained with primary antibodies (CD4 AF 647, CD8α, CD45.2 AF 488) diluted in 1% BSA in PBS at room temperature for 1 h, followed by staining with the secondary antibody (goat anti-rabbit, AF 555) for 1 h and nuclear staining with DAPI at room temperature for 15 min. Samples were mounted with Prolong Gold antifade mountant (Invitrogen #P36930) and images were acquired on confocal microscope Leica TCS SP8 (Leica Microsystems, objective HC PL APO 40x/1.30 OIL CS2).

### Production and testing of recombinant GITRL

The DNA coding sequences of tagged recombinant GITRL and OX40L were produced by GeneArt Gene Synthesis service (ThermoFisher Scientific). The sequences of GITRL and OX40L constructs contain: human CD33 signal peptide (AA 1-17, Uniprot: P20138), 6× His, 2× Strep, 1× Flag tag, and murine 47-173 AA of murine GITRL (Uniprot: Q7TS55) or 87-197 AA OX40L (Uniprot: P43488). The constructs were inserted in pcDNA3.1 vector for subsequent production in HEK293FT cells. HEK293FT cells were transfected with the plasmid (30 µg) using polyethylenimine (75 µg) transfection. Supernatants were collected after 3 days post transfection and purified using GraviTrap TALON columns (GE Healthcare # GE29-0005-94). The columns were first equilibrated with purification buffer (50 mM sodium phosphate pH 7.4, 300 mM NaCl) before sample loading. Following, the columns were washed with 20 mM imidazole in purification buffer and eluted using 350 mM imidazole in purification buffer. Samples were then loaded on Amicon Ultra-15 centrifugal filters (10 kDa molecular weight cutoff, Merck Millipore # UFC9010) for imidazole removal and washed with purification buffer. Samples were concentrated by centrifugation and mixed with glycerol 1:1 for long-term storage in -80°C. To test the production, GITRL and OX40L were mixed with Laemli sample buffer +/- 50 mM dithiothreitol (DTT, Sigma #D0632) and tested using WB and visualization with InstantBlue Coomassie protein stain (Expedeon #ISB1L). To confirm the ability of ligands to bind the receptor on T cells, non-activated and PMA/Iono pre-activated cells were treated with GITRL or OX40L in FACS buffer (PBS/2% FBS/0.1% azide). The binding of GITRL or OX40L was detected using anti-FLAG APC antibody and measured on Accuri C6 cytometer (BD Bioscience). The data were analyzed using FlowJo software (BD Bioscience).

### Primary cells activation for immunoprecipitation

T cells were isolated from C57Bl/6J mice and incubated with PMA (5 ng/ml) and Ionomycin (0.2 µM) in complete RPMI (10% FBS/100 U/ml penicillin/100 µg/ml streptomycin/40 µg/ml gentamicin) for 3 days. Following, cell suspension was split for negative control, ‘post-lysis’ control, and the sample. Samples were incubated for 30 min at 37°C in serum-free media and activated with GITRL or OX40L (500 ng/ml, the sample only). After the indicated time cells were lysed in 1% *n*-Dodecyl-β-D-Maltoside (DDM, Thermo Scientific #89903) in lysis buffer (30 mM Tris pH 7.4, 120 mM NaCl, 2mM KCl, 2 mM EDTA, 10% glycerol, 10 mM chloroacetamide (Sigma #C0267), 10 mM cOmplete protease inhibitor cocktail (Roche #5056489001), and PhosSTOP tablets (Roche #4906837001)). Samples and controls were incubated at 4°C for 30 min and cleared by centrifugation for 30 min, 2°C, 21,130 g. Samples were used for following immunoprecipitation or mixed with Laemli sample buffer, reduced by 50 mM DTT, heated for 3 min at 92°C, and analyzed by immunoblotting.

### Immunoprecipitation of receptors proximal signaling complex

After the activation step, to the ‘post-lysis’ control, 1.6 µg of GITRL or OX40L was added to the cell lysate. Supernatant was collected and mixed with the anti-FLAG M2 beads and incubated overnight at 4°C. Beads were then washed three times with 10× diluted lysis buffer containing 0.1% of DDM and eluted with Laemli sample buffer, reduced by 50 mM DTT, heated for 3 min at 92°C, and analyzed by immunoblotting.

### Tandem affinity purification of receptor proximal signaling complexes for MS

After the activation step, to the ‘post-lysis’ sample, 3 µg of GITRL, or OX40L were added. For the first purification step, samples were mixed with anti-FLAG M2 beads (Sigma #A2220) and incubated at 4°C overnight. Samples were then washed with lysis buffer containing 0.1% DDM and incubated with lysis buffer containing 1% DDM and 3×Flag peptide (100 µg/ml, Sigma #F4799) overnight for elution. Subsequently, the supernatant was collected and the elution step was repeated with 8h incubation. The second purification step was done with 50 µl of Strep-Tactin beads (IBA Life-science #2-1201-010) overnight. After incubation, samples were washed 3× with lysis buffer containing 0.1% DDM and 1× with lysis buffer only. Elution step was done with MS Elution buffer (2% sodium deoxycholate in 50 mM Tris pH 8.5).

### Protein digestion and MS analysis

The eluted protein samples (200 µl) were reduced with 5 mM tris(2-carboxyethyl)phosphine at 60°C for 60 minutes and alkylated with 10 mM methyl methanethiosulfonate at room temperature for 10 min. Proteins were cleaved overnight with 1 µg of trypsin (Promega) at 37°C. To remove sodium deoxycholate, samples were acidified with 1% trifluoroacetic acid, mixed with an equal volume of ethyl acetate, centrifuged (15,700 g, 2 min), and an aqueous phase containing peptides was collected [45]. This step was repeated two more times. Peptides were desalted using in-house-made stage tips packed with C18 disks (Empore) [46] and resuspended in 20 µl of 2% acetonitrile with 1% trifluoroacetic acid.

The digested protein samples (12 µl) were loaded onto the trap column (Acclaim PepMap300, C18, 5 µm, 300 Å Wide Pore, 300 µm x 5 mm) using 2% acetonitrile with 0.1% trifluoroacetic acid at a flow rate of 15 μl/min for 4 min. Subsequently, peptides were separated on a Nano Reversed-phase column (EASY-Spray column, 50 cm x 75 µm internal diameter, packed with PepMap C18, 2 µm particles, 100 Å pore size) using a linear gradient from 4% to 35% acetonitrile containing 0.1% formic acid at a flow rate of 300 nl/min for 60 minutes.

Ionized peptides were analyzed on a Thermo Orbitrap Fusion (Q-OT-qIT, Thermo Scientific). Survey scans of peptide precursors from 350 to 1400 m/z were performed at 120K resolution settings with a 4×10^5^ ion count target. Four different types of tandem MS were performed according to precursor intensity. The first three types were detected in Ion trap in rapid mode, and the last one was detected in Orbitrap with 15000 resolution settings: 1) For precursors with intensity between 1×10^3^ to 7×10^3^ with CID fragmentation (35% collision energy) and 250 ms of ion injection time. 2) For ions with intensity in the range from 7×10^3^ to 9×10^4^ with CID fragmentation (35% collision energy) and 100 ms of ion injection time. 3) For ions with intensity in the range from 9×10^4^ to 5×10^6^ with HCD fragmentation (30% collision energy) and 100 ms of ion injection time. 4) For intensities 5×10^6^ and more with HCD fragmentation (30% collision energy) and 35 ms of ion injection time. The dynamic exclusion duration was set to 60 s with a 10 ppm tolerance around the selected precursor and its isotopes. Monoisotopic precursor selection was turned on. The instrument was run in top speed mode with 3 s cycles.

All MS data were analyzed and quantified with the MaxQuant software (version 1.6.15.0) [47]. The false discovery rate (FDR) was set to 1% for both proteins and peptides and the minimum length was specified as seven amino acids. The Andromeda search engine was used for the MS/MS spectra search against the murine Swiss-Prot database (downloaded from Uniprot in December 2020). Trypsin specificity was set as C-terminal to Arg and Lys, also allowing the cleavage at proline bonds and a maximum of two missed cleavages. β-methylthiolation, N-terminal protein acetylation, carbamidomethylation, Met oxidation, and STY phosphorylation were included as variable modifications. Label-free quantification was performed using the Intensity Based Absolute Quantification (iBAQ) algorithm, which divides the sum of all precursor-peptide intensities by the number of theoretically observable peptides [48]. Data analysis was performed using Perseus 1.6.14.0 software [49].

### T2-Kb OVA loading

T2-Kb cells were stained in RPMI with DDAO dye (2.5µM) for 15 min at 37°C, 5% CO_2_. Following, cells were resuspended in RPMI (10% FBS, 100 U/ml penicillin, 100 mg/ml streptomycin, 40 mg/ml gentamicin) and split into 48-well plate and loaded with indicated concentration of OVA peptide for 2 h at 37°C, 5% CO_2_.

### Primary T-cell activation

Primary T cells were isolated from the lymph nodes and/or spleen. The spleen was incubated in ACK buffer for 3 min in room temperature to remove red blood cells.

For the activation of T cells using T2-Kb cells, CD8^+^ T cells were enriched using magnetic separation (MACS). Following, T cells were incubated for 30 min in serum-free RPMI media. Cells were incubated for 2 h with OVA-loaded T2-Kb cells in 2:1 ratio. After activation, cells were fixed with formaldehyde (2%, Sigma #F8775) and permeabilized with 0.3% Triton X-100 in FACS buffer (PBS, 5% goat serum, 2mM EDTA, 0.1% sodium azide), followed by overnight staining with specific primary antibodies at room temperature. Next, samples were stained with a fluorescently labeled secondary antibody for 45 min and analyzed by flow cytometry.

To study proliferation of T cells, cells were stained with Cell Trace Violet (1000× diluted) for 10 min at 37°C, 5% CO_2_. Cells were resuspended IMDM (10% FBS, 100 U/ml penicillin, 100 mg/ml streptomycin, 40 mg/ml gentamicin) and split to 48-well plate coated with indicated concentration of anti-CD3ε or Kb-OVA monomer. Cells were incubated for 72 h at 37°C, 5% CO_2_ and analyzed by flow cytometry.

For the analysis of expression of effector molecules, cells were (i) left untreated, (ii) treated with p38 inhibitor (12.5 µM), (iii) activated with anti-CD3/CD28 beads at cell to cells ratio 1:1, or (iv) activated with anti-CD3/CD28 beads an beads to cells ratio 1:1 and treated with p38 inhibitor (12.5 µM). The incubation time was 16 h at 37°C, 5% CO_2_. After 12 h, Golgi stop was added (500×) for 4 h. Samples were fixed and permeabilized with BD Cytofix/Cytoperm Fixation/Permeabilization Kit was used (BD Bioscience #554714) according to the manufacturer’s protocol. Samples were stained with indicated antibodies and analyzed by flow cytometry.

For the analysis of surface activation markers, CD8^+^ cells were FACS sorted from the lymph nodes. Cells were activated with anti-CD3ε/CD28 (ThermoFisher #11453D) beads at indicated beads to cell ratios for 16 hours. Samples were then stained with indicated antibodies and analyzed by flow cytometry.

For the analysis of GITR signaling, cells were activated with PMA (5 ng/ml) and ionomycin (0.5 µM) overnight. Following, cells were incubated in IMDM (10% FBS, 100 U/ml penicillin, 100 mg/ml streptomycin, 40 mg/ml gentamicin) supplemented with IL-2 for 72 h. Next, cells were activated with recombinant GITRL (500 ng/ml) for 15 min at 37°C. Cells were fixed with 2% formaldehyde and permeabilized using 90% methanol. Samples were stained with indicated primary antibodies overnight at room temperature. Following, samples were stained with indicated fluorescently labeled secondary antibodies and analyzed by flow cytometry.

### In vitro suppression assay

CD4^+^ and CD8^+^ T cells were FACS sorted from C57Bl/6J WT mice and labeled with the CTV dye. Tregs were FACS sorted based on CD4^+^GFP^+^ expression from ABIN1 WT or GTKO DEREG mice. 48-well tissue culture plate was coated with anti-CD3ε antibody. Conventional cells were plated to 48-well plate in complete IMDM (10% FBS, 100 U/ml penicillin, 100 mg/ml streptomycin, 40 mg/ml gentamicin) in 1:1 ratio with Tregs, one well with no Tregs was used as control. Cells were incubated at 37°C, 5% CO_2_ for 72 h and measured by flow cytometry.

### Bulk RNAseq

1×10^6^ CD8^+^ cells were FACS sorted from lymph nodes and spleen of OT-I ABIN1 WT or GTKO littermates per sample. Non-activated controls were directly used for RNA isolation. The remaining cells were activated with anti-CD3ε/CD28 (ThermoFisher #11453D) beads at indicated beads to cell ratios for 16 hours prior to the RNA isolation. In some experiments, the samples were treated with p38 inhibitor SB203580 (12.5 µM) for the whole activation period. The RNA was isolated using RNA Clean & Concentrator-5 kit (Zymoresearch #R1014) according to the manufacturer’s instruction. The cDNA libraries were prepared using KAPA mRNA Hyperprep Kit (Roche #KK8580) according to the manufacturer’s instructions. The single-end sequencing was performed on Illumina NextSeq 500 using NextSeq 500/550 High Output Kit v2.5 (75 cycles) (Illumina #20024906) with final reads having length of 76bp. The base calling was performed using Illumina BaseSpace GenerateFASTQ workflow v1.37.0.

### Data analysis

The quality of reads was verified using FastQC 0.11.9 and MultiQC 1.13 tools [50, 51]. The reads were then aligned against mouse genome GRCm39 obtained from Ensembl v106 [52] using STAR aligner 2.7.10a [53]. The analysis was done on R 4.2.1. The quantification of results was done using R packages tximport 1.24.0 and GenomicAlignments 1.32.1 [54–56]. The genes in count matrices were annotated using AnnotationHub 3.4.0 [57] using the same version of Ensembl database as for the alignment. The downstream analysis including normalization, raw count modelling and differential expression analyses on complete count matrices as well as subsets of specific samples was done using R package DESeq2 1.36.0 [56]. For differential expression analysis, the log2-fold changes were shrunk using DESeq2 option “ashr” [58]. The visualization of data was done using from normalized counts transformed by unblinded log regularization. The heatmaps has been corrected for batch effects using package limma 3.52.2 [59]. The genes upregulated in the *Abin1*^GTKO/GTKO^ cells were selected as genes with fold-change over 1.5 in the activated cells from the *Abin1*^GTKO/GTKO^ mice vs. the activated cells from the *Abin1*^WT/WT^ mice (both without the p38 inhibitor) in each of the two experiments that were performed. The GSEA was then performed on these selected genes using R package fgsea 1.22.0 [60]. The genes upregulated in the ABIN1^GTKO/GTKO^ cells were selected as genes with the average upregulation in the cells from the ABIN1^GTKO/GTKO^ mice compared to the cells from the ABIN1^WT/WT^ mice in each of the two experiments that were performed. The GSEA was then performed on these selected genes using R package fgsea 1.22.0 [61].

### Autoimmune diabetes

The model of autoimmune diabetes was described previously [22]. Briefly, bone marrow-derived dendritic cells (BMDCs) derived from the bone marrow of congenic Ly5.1 C57BL/6J mice were cultured in IMDM media supplemented with 10% FBS (GIBCO), 100 U/ml penicillin (BB Pharma), 100 mg/ml streptomycin (Sigma-Aldrich), 40 mg/ml gentamicin (Sandoz), and with 2% of supernatant from J558 cells producing GM-CSF for 10 days at 37°C, 5% CO_2_ for 10 days [62] and finally matured with LPS (25 µg) and loaded with 50 µg OVA peptide (SIINFEKL) for 3 h. Indicated numbers of T cells isolated from ABIN1^WT/WT^ or ABIN1^GTKO/GTKO^ OT-I Rag2^KO/KO^ mice were adoptively transferred to RIP.OVA mice intravenously. On the following day, 1×10^6^ OVA-loaded BMDCs were injected intravenously. The glucose in urine was monitored with test strips on daily basis (GLUKOPHAN, Erba Lachema) for 2 weeks. The blood glucose was measured using Contour blood glucose meter (Bayer) on day 7 post immunization. The animal was considered as diabetic when the concentration of glucose in the urine was higher than 1000 mg/dl for two consecutive days.

### The tumor infiltration model

5×10^5^ MC-38 cells expressing OVA [63] were injected subcutaneously to Ly5.1/Ly5.2 heterozygous mice. After nine days, OT-I cells were isolated from Ly5.1 mouse and mixed with Ly5.2 OT-I T cells isolated from the ABIN1^GTKO/GTKO^ mice or their ABIN1^WT/WT^ littermates at 1:1 ratio. Total number of 2×10^6^ OT-I cells was injected i.v. One week after OT-I injection, cells from tumor, dLN, ndLN and spleen were isolated and measured by flow cytometry. Tumors were cut into small pieces and incubated with Liberase (100 µg/ml, Roche #5401020001) and DNAse I (50 µg/ml, Roche #101104159001) in wash buffer (1% BSA and 1 mM EDTA in HBSS without Ca^2+^/Mg^2+^) at 37°C, 350 RPM shaking for 45 min. Every 10 min, the samples were resuspended with wide-bore pipette tip (1 ml). After the enzymatic digestion, samples were filtered through 100 µm strainer and centrifuged 350g at 4°C for 3 min. The pellet was resuspended in 5 ml of 40% Percoll (Cytiva, 17089101) in DMEM. To the bottom of the tube, 5 ml of 80% Percoll was added to create a gradient. Samples were spin 320g, at room temperature acc/dec 0, for 23 min. The interphase that contained the lymphocytes was collected and centrifuged 400g, 4°C for 5 min. Following, the samples were analyzed by flow cytometry.

### The assessment of the anti-tumor response

5×10^5^ MC-38 cells expressing OVA were injected subcutaneously to the CD3ε^KO/KO^ host mice. Indicated numbers of OT-I T cells from ABIN1^WT/WT^ or ABIN1^GTKO/GTKO^ were adoptively transferred intravenously. Tumor growth was monitored till day 23 after OT-I injection using caliper. Tumor volume was calculated using the formula V = (L × S^2^)/2 (L is the longest diameter, S is the shortest diameter). When the tumor reached the endpoint volume of 500 mm^3^, the mouse was sacrificed.

### Listeria and LCMV

Ly5.1 or Ly5.1/Ly5.2 heterozygotes were used as host mice. At day -1, sorted OT-I cells from ABIN1^WT/WT^ or ABIN1^GTKO/GTKO^ mice were adoptively transferred, 5×10^4^ or 10×10^4^ per mouse depending on experimental setup. Following, at day 0 host mice were infected intravenously with 5000 CFU of Lm-OVA or Lm-Q4H7 or 2-3 × 10^5^ PFU of LCMV-OVA intraperitonealy, respectively. On day 6 (Lm) or 5 (LCMV), splenocytes were analyzed by flow cytometry.

### Imaging cytometry analysis

T cells from ABIN1^WT/WT^ or ABIN1^GTKO/GTKO^ OT-I Rag2^KO/KO^ mice were FACS sorted and activated in vitro with OVA-loaded T2-Kb cells for 1 h. Following activation, cells were fixed with 2% formaldehyde and permeabilized using 0.3% Triton-X in PBS. Samples were stained with indicated primary antibodies at room temperature overnight. Next day, samples were stained with goat anti-rabbit secondary antibody conjugated with AF 488 for 45 min and measured on Amnis Imagestream X Mk II Imaging Flow Cytometer (Luminex). Data were analyzed using Ideas 6.2 software (Amnis).

### Bone marrow chimeras

Bone marrow cells were isolated from femurs of Ly5.1/Ly5.2 ABIN1^WT/WT^ or Ly5.2 ABIN1^GTKO/GTKO^ mice. Cells were mixed in 1:1 ratio and total number of 10^6^ cells was injected into lethally irradiated (6 Gy, X-RAD 225 XL) congenic Ly5.1 mice intravenously. The splenocytes and lymph node cells of these mice were analyzed after 8 weeks by flow cytometry.

### Data and code availability

The raw sequences as well as generated raw count matrices are available at GEO NCBI under accession number GSE245397. The code used to generate counts, downstream analyses and figures for this publication are available at https://github.com/Lab-of-Adaptive-Immunity/ABIN_KO_project.

