## Supplementary material for "ABIN1 is a negative regulator of effector functions in cytotoxic T cells": Fig. EV

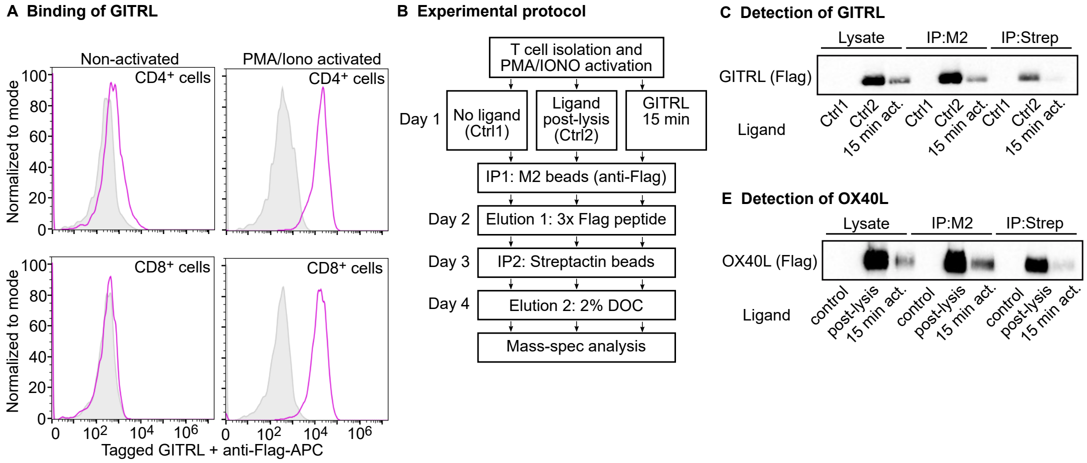


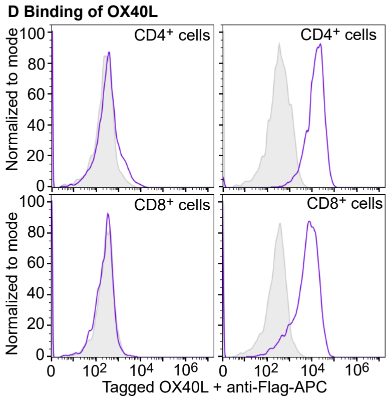


**F OX40-SC analysis: Number of identified peptides**

| | No ligand | OX40L post-lysis | OX40L 15 min |
|---|---|---|---|
| OX40L | - | 11 | 11 |
| OX40 | - | 12 | 7 |
| TRAF1 | - | - | 6 |
| TRAF2 | - | - | 5 |
| TRAF3 | - | - | 8 |
| ABIN1 | - | - | 7 |
| A20 | - | - | 15 |

**G OX40-SC analysis: Coverage (%)**

| | No ligand | OX40L post-lysis | OX40L 15 min |
|---|---|---|---|
| OX40L | - | 41 | 41 |
| OX40 | - | 40 | 31 |
| TRAF1 | - | - | 21 |
| TRAF2 | - | - | 14 |
| TRAF3 | - | - | 18 |
| ABIN1 | - | - | 18 |
| A20 | - | - | 27 |

**Extended View Figure 1. Analysis of the proximal GITR and OX40 signaling complex (SC). (A)** Primary murine T cells were pre-activated with PMA/ionomycine for 72 hours or not and stained with the anti-CD4 and anti-CD8 antibodies, recombinant GITRL followed by anti-FLAG antibody. Representative experiments out of 3 in total are shown. **(B)** Illustration of the protocol for the identification of the composition of GITR and OX40 signaling complexes. **(C)** Detection of the recombinant GITRL in the lysate and after the first and second affinity purification step by immunoblotting. A representative experiments out of 3 in total. **(D)** Primary murine T cells were pre-activated with PMA/ionomycine for 72 hours or not and stained with the anti-CD4 and anti-CD8 antibodies, recombinant OX40L followed by anti-FLAG antibody. Representative experiments out of 3 in total are shown. **(E)** Detection of the recombinant GITRL in the lysate and after the first and second affinity purification step by immunoblotting. A representative experiments out of 3 in total. **(F-G)** Primary murine T cells mice were pre-activated with PMA/ionomycin for 72 hours and stimulated with the recombinant OX40L for 15 min. The cells were lysed and the OX40-SC was isolated via tandem affinity purification and samples were analyzed by mass spectrometry. Results of the protein identification are shown as the number of peptides (G) and the coverage (H). A single experiment was performed.

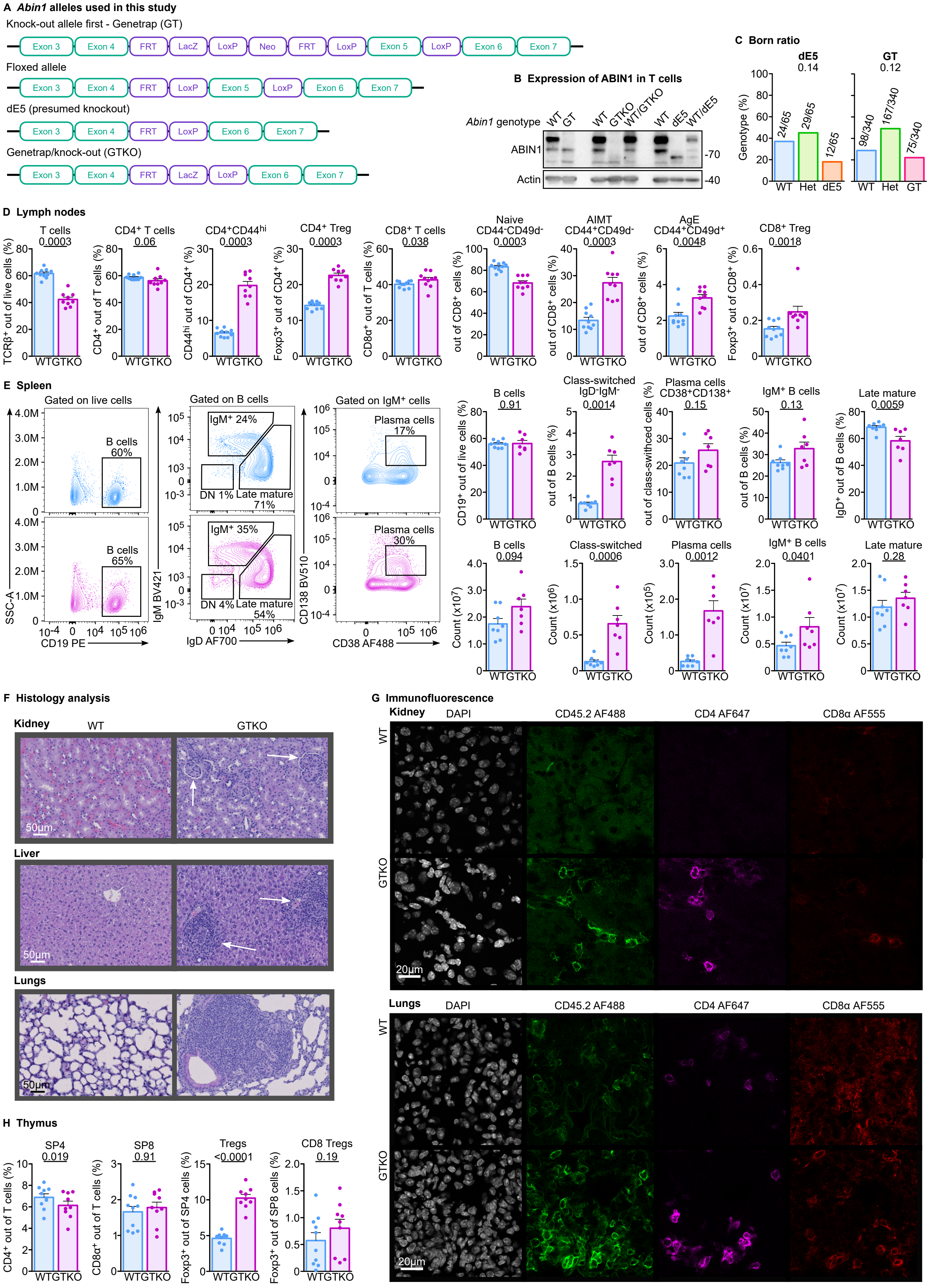
A Abin1 alleles used in this study
Knock-out allele first - Genetrap (GT)
Exon 3 Exon 4 FRT LacZ LoxP Neo FRT LoxP Exon 5 LoxP Exon 6 Exon 7
Floxed allele
Exon 3 Exon 4 FRT LoxP Exon 5 LoxP Exon 6 Exon 7
dE5 (presumed knockout)
Exon 3 Exon 4 FRT LoxP Exon 6 Exon 7
Genetrap/knock-out (GTKO)
Exon 3 Exon 4 FRT LacZ LoxP Exon 6 Exon 7
B Expression of ABIN1 in T cells
Abin1 genotype WT GT WT GTKO WT/GTKO WT dE5 WT/dE5
ABIN1
Actin
-70
-40
C Born ratio
dE5 0.14
GT 0.12
Genotype (%)
24/65 29/65 12/65
98/340 167/340 75/340
WT Het dE5
WT Het GT
D Lymph nodes
T cells 0.0003
CD4+ T cells 0.06
CD4+CD44hi 0.0003
CD4+ Treg 0.0003
CD8+ T cells 0.038
Naive CD44-CD49d- 0.0003
AIMT CD44+CD49d- 0.0003
AgE CD44+CD49d+ 0.0048
CD8+ Treg 0.0018
TCRβ+ out of live cells (%)
CD4+ out of T cells (%)
CD44hi out of CD4+ (%)
Foxp3+ out of CD4+ (%)
CD8α+ out of T cells (%)
out of CD8+ cells (%)
Foxp3+ out of CD8+ (%)
WT GTKO
E Spleen
Gated on live cells
Gated on B cells
Gated on IgM+ cells
B cells 60%
B cells 65%
IgM+ 24%
IgM+ 35%
DN 1% Late mature 71%
DN 4% Late mature 54%
Plasma cells 17%
Plasma cells 30%
SSC-A
CD19 PE
IgM BV421
IgD AF700
CD138 BV510
CD38 AF488
B cells 0.91
Class-switched IgD-IgM- 0.0014
Plasma cells CD38+CD138+ 0.15
IgM+ B cells 0.13
Late mature 0.0059
CD19+ out of live cells (%)
out of B cells (%)
out of class-swithced cells (%)
IgD+ out of B cells (%)
B cells 0.094
Class-switched 0.0006
Plasma cells 0.0012
IgM+ B cells 0.0401
Late mature 0.28
Count (x10^7)
Count (x10^6)
Count (x10^5)
F Histology analysis
Kidney WT GTKO
Liver
Lungs
50µm
G Immunofluorescence
Kidney DAPI CD45.2 AF488 CD4 AF647 CD8α AF555
Lungs DAPI CD45.2 AF488 CD4 AF647 CD8α AF555
WT
GTKO
20µm
H Thymus
SP4 0.019
SP8 0.91
Tregs <0.0001
CD8 Tregs 0.19
CD4+ out of T cells (%)
CD8α+ out of T cells (%)
Foxp3+ out of SP4 cells (%)
Foxp3+ out of SP8 cells (%)
WT GTKO

**Extended View Figure 2. Characterization of Abin1$^{GT/GT}$ (GT), Abin1$^{dE5/dE5}$ (dE5), and Abin1$^{GTKO/GTKO}$ (GTKO) mice. (A)** An overview of *Abin1* alleles used in this study. **(B)** Immunoblot analysis ABIN1 in GT, GTKO, dE5 mice and corresponding littermate controls. A representative experiment is shown, GT in 5 independent experiments, GTKO in 3 independent experiments, dE5 in 2 independent experiments. **(C)** Heterozygous Abin1$^{WT/dE5}$ or Abin1$^{WT/GT}$, respectively, were bred and the genotype of the offspring was determined upon weaning. The frequencies and numbers of pups with particular genotypes are indicated. n = 340 (GT) or 65 (dE5) offspring mice in total from 16 (GT) or 2 (dE5) breedings. **(D)** Lymph node cells were stained with indicated antibodies and analyzed by flow cytometry. Aggregate results of the abundance of indicated subsets are shown, n = 10 (WT) or 9 (GTKO) mice in 6 independent experiments. **(E)** Splenocytes were stained with indicated antibodies and analyzed by flow cytometry. Representative dot plots and aggregate results of the frequency of indicated subsets are shown. n = 10 (WT) or 9 (GTKO) mice in 6 independent experiments. **(F)** Histological analysis using H&E staining of indicated organs of 20-26 week old mice. Arrows pointing to inflammatory foci. Representative staining out of 4 mice per group in total. **(G)** Cryosections of lungs and kidneys of WT and GTKO mice were stained with indicated antibodies and DAPI (nuclei) and analyzed by confocal fluorescence microscopy. Representative sections out of 4 mice per group in total. **(H)** Fixed and permeabilized thymocytes from WT and GTKO mice were stained with indicated antibodies and analyzed by flow cytometry. Aggregate results of the abundance of indicated subsets are shown. n = 10 (WT) or 9 (GTKO) in 6 independent experiments. When applicable, the results are shown as means + SEM and p-values are indicated. Statistical significance was determined by a binomial test (A) or two-tailed Mann-Whitney test (D, E, H).

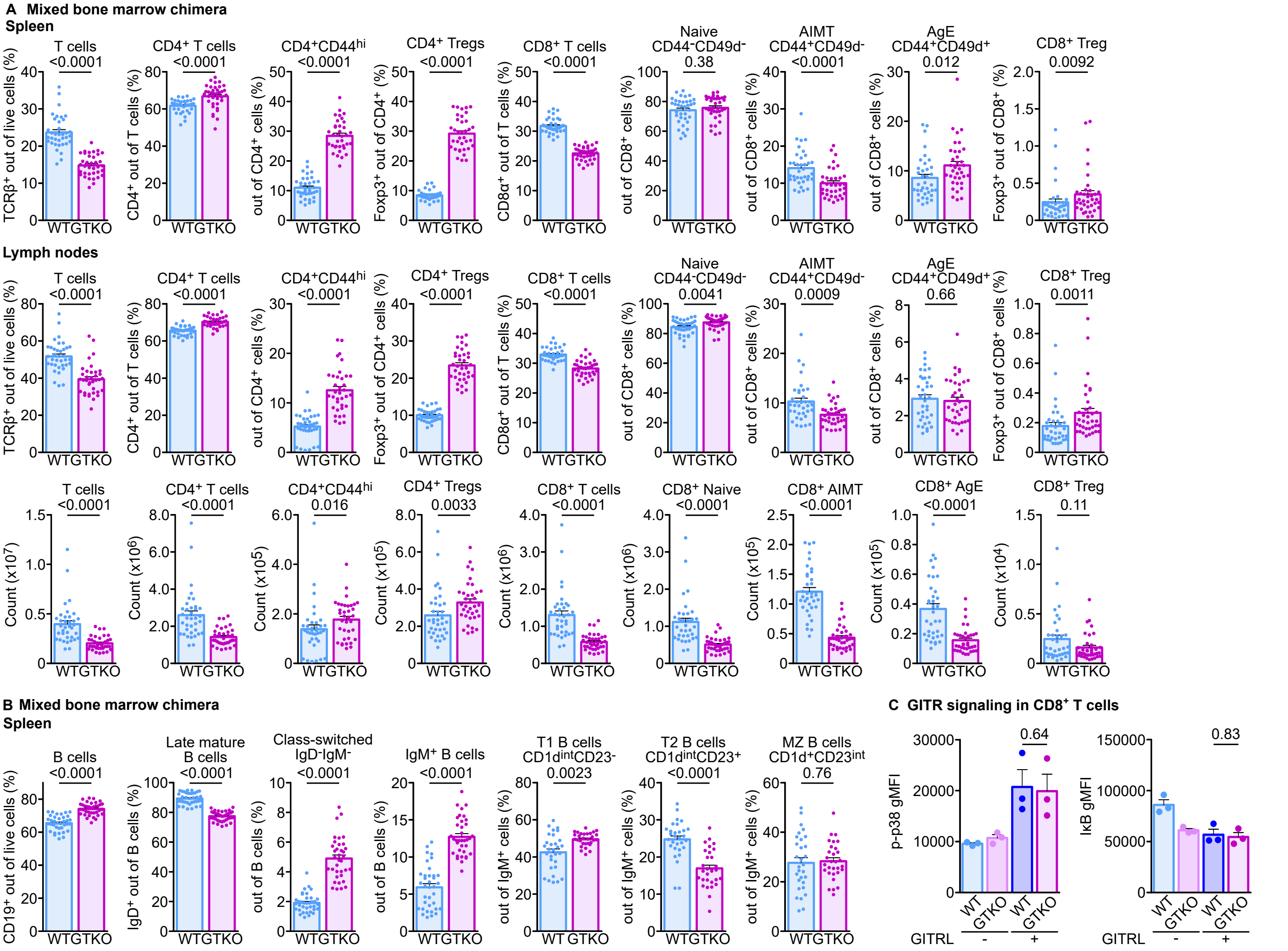
A Mixed bone marrow chimera
Spleen
T cells <0.0001
CD4+ T cells <0.0001
CD4+CD44hi <0.0001
CD4+ Tregs <0.0001
CD8+ T cells <0.0001
Naive CD44-CD49d- 0.38
AIMT CD44+CD49d- <0.0001
AgE CD44+CD49d+ 0.012
CD8+ Treg 0.0092
Lymph nodes
T cells <0.0001
CD4+ T cells <0.0001
CD4+CD44hi <0.0001
CD4+ Tregs <0.0001
CD8+ T cells <0.0001
Naive CD44-CD49d- 0.0041
AIMT CD44+CD49d- 0.0009
AgE CD44+CD49d+ 0.66
CD8+ Treg 0.0011
T cells <0.0001
CD4+ T cells <0.0001
CD4+CD44hi 0.016
CD4+ Tregs 0.0033
CD8+ T cells <0.0001
CD8+ Naive <0.0001
CD8+ AIMT <0.0001
CD8+ AgE <0.0001
CD8+ Treg 0.11
B Mixed bone marrow chimera
Spleen
B cells <0.0001
Late mature B cells <0.0001
Class-switched IgD-IgM- <0.0001
IgM+ B cells <0.0001
T1 B cells CD1dintCD23- 0.0023
T2 B cells CD1dintCD23+ <0.0001
MZ B cells CD1d+CD23int 0.76
C GITR signaling in CD8+ T cells
p-p38 gMFI
IkB gMFI
0.64
0.83
GITRL - +
WT GTKO

**Extended View Figure 3**. **Intrinsic roles of ABIN1 in T cells. (A, B)** The experiment shown in Fig. 3A-B. Aggregate results of the frequency and absolute counts of indicated subsets of T cells (A) and B cells (B) are shown. n = 36 mice in 5 independent experiments. **(C)** The experiment shown in Figure 3D. Lymph node cells from *Abin1*$^{WT/WT}$ or *Abin1*$^{GTKO/GTKO}$ mice were pre-activated with PMA/ionomycin and stimulated with GITRL or left untreated (controls). Indicated activation pathways were analyzed by flow cytometry. Aggregate results of phospho-p38 and IκB levels in $CD8^+$ T cells. n = 4 independent experiments. The results are shown as means + SEM and p-values are indicated. Statistical significance was determined by two-tailed Mann-Whitney test (A-B) or two-tailed Student's t test (C).

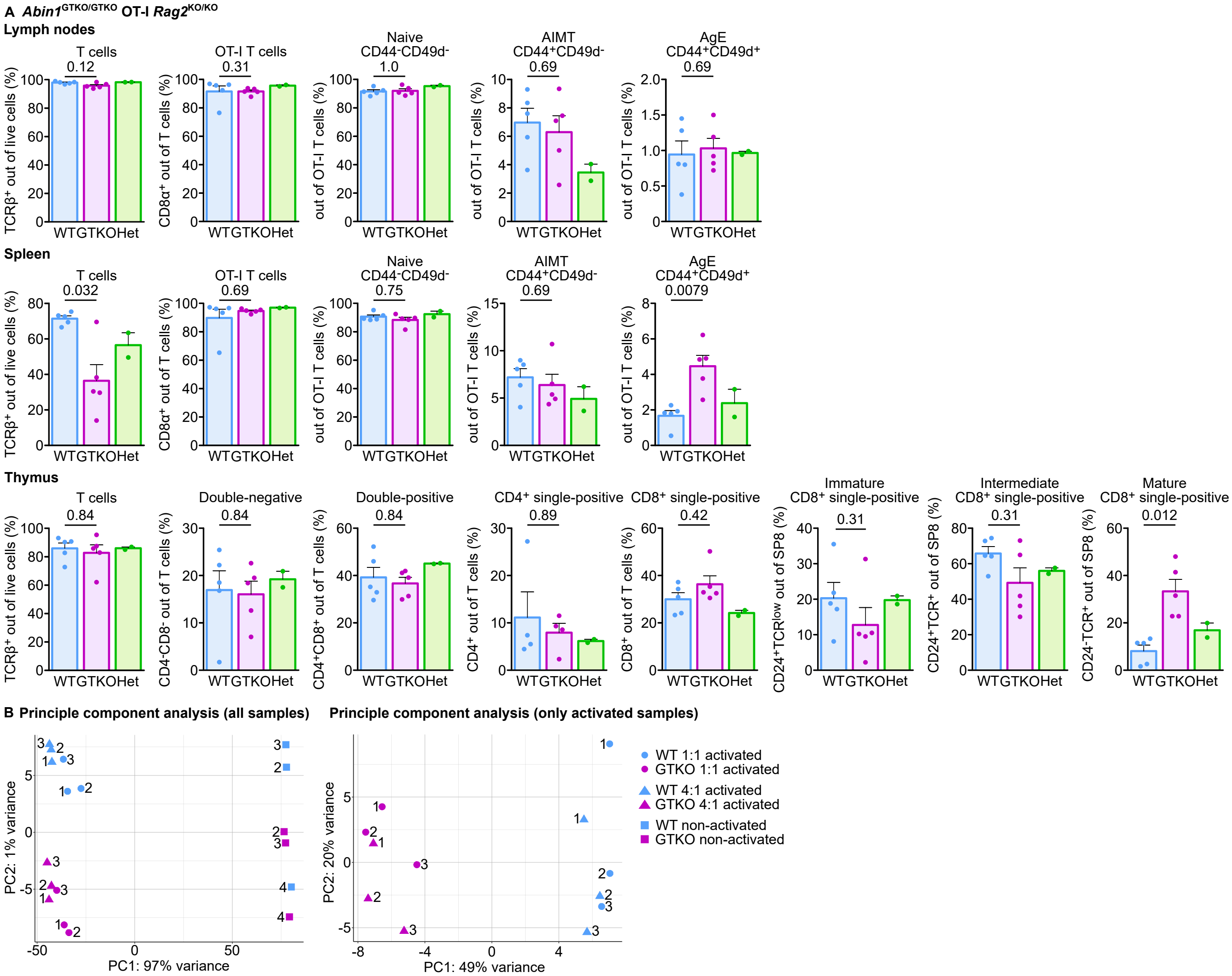
A Abin1GTKO/GTKO OT-I Rag2KO/KO
Lymph nodes
T cells
OT-I T cells
Naive CD44-CD49d-
AIMT CD44+CD49d-
AgE CD44+CD49d+
0.12
0.31
1.0
0.69
0.69
TCRβ+ out of live cells (%)
CD8α+ out of T cells (%)
out of OT-I T cells (%)
WT GTKO Het
Spleen
0.032
0.69
0.75
0.69
0.0079
Thymus
Double-negative
Double-positive
CD4+ single-positive
CD8+ single-positive
Immature CD8+ single-positive
Intermediate CD8+ single-positive
Mature CD8+ single-positive
0.84
0.84
0.84
0.89
0.42
0.31
0.31
0.012
CD4-CD8- out of T cells (%)
CD4+CD8+ out of T cells (%)
CD4+ out of T cells (%)
CD8+ out of T cells (%)
CD24+TCRlow out of SP8 (%)
CD24+TCR+ out of SP8 (%)
CD24-TCR+ out of SP8 (%)
B Principle component analysis (all samples)
Principle component analysis (only activated samples)
PC1: 97% variance
PC2: 1% variance
PC1: 49% variance
PC2: 20% variance
WT 1:1 activated
GTKO 1:1 activated
WT 4:1 activated
GTKO 4:1 activated
WT non-activated
GTKO non-activated

**Extended View Figure 4. Characterization of *Abin1*$^{GTKO/GTKO}$ OT-I mice. (A)** Cells from lymph nodes, spleen, and thymus from *Abin1*$^{WT/WT}$ OT-I *Rag2*$^{KO/KO}$ (WT), *Abin1*$^{GTKO/GTKO}$ OT-I *Rag2*$^{KO/KO}$ (GTKO), and *Abin1*$^{WT/GTKO}$ OT-I *Rag2*$^{KO/KO}$ (HET) were stained with indicated antibodies and analyzed by flow cytometry. Aggregate results of the frequency of indicated subsets are shown. n = 5 (WT and GTKO) or 2 (WT/GTKO) mice in 5 independent experiments. **(B)** Principal component analysis of the RNAseq experiment shown in Figure 4C-D.

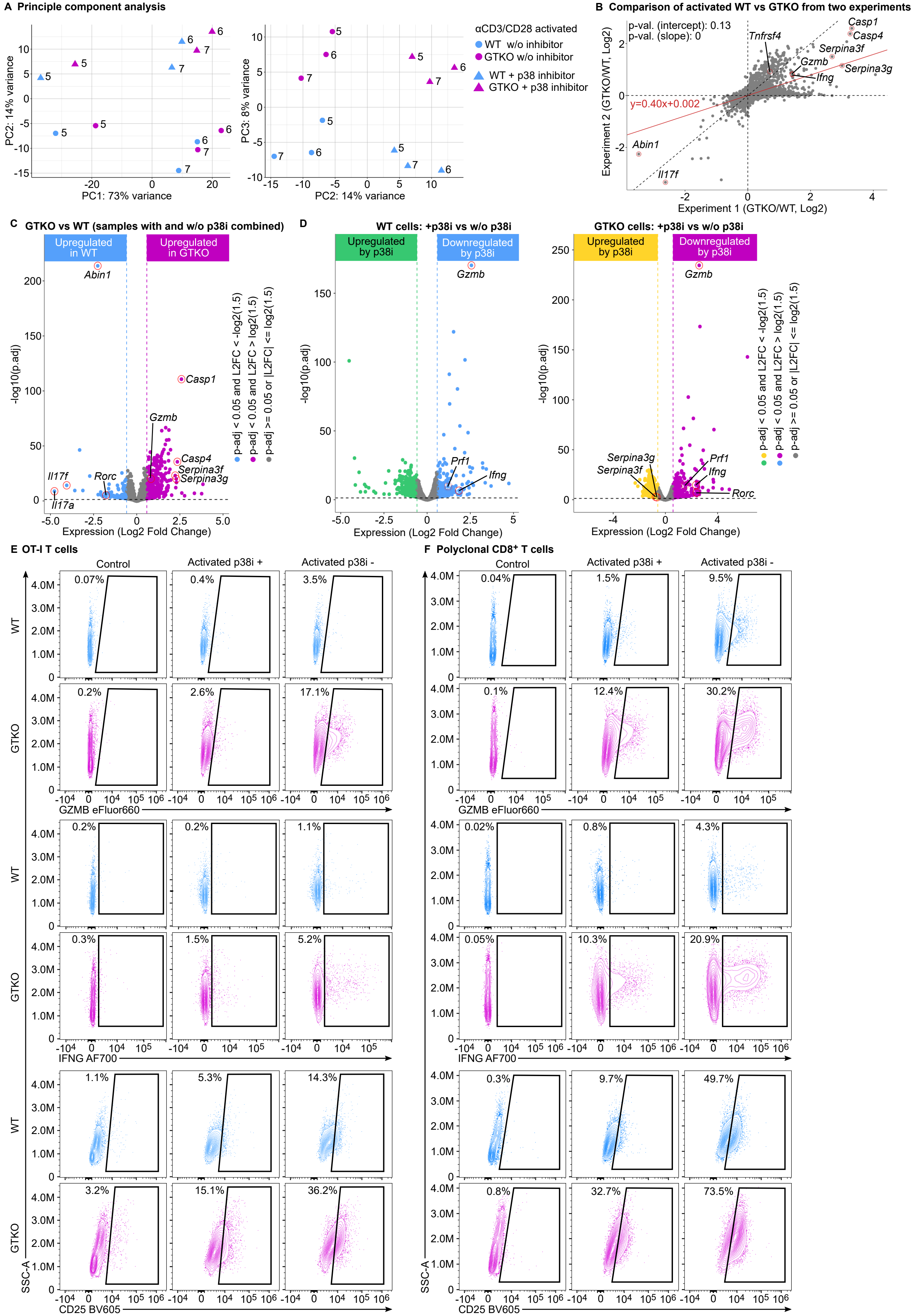

A Principle component analysis
PC1: 73% variance
PC2: 14% variance
PC3: 8% variance
αCD3/CD28 activated
WT w/o inhibitor
GTKO w/o inhibitor
WT + p38 inhibitor
GTKO + p38 inhibitor
B Comparison of activated WT vs GTKO from two experiments
p-val. (intercept): 0.13
p-val. (slope): 0
y=0.40x+0.002
Experiment 1 (GTKO/WT, Log2)
Experiment 2 (GTKO/WT, Log2)
Casp1
Casp4
Tnfrsf4
Serpina3f
Gzmb
Serpina3g
Ifng
Abin1
Il17f
C GTKO vs WT (samples with and w/o p38i combined)
Upregulated in WT
Upregulated in GTKO
Abin1
Casp1
Gzmb
Casp4
Serpina3f
Serpina3g
Il17f
Rorc
Il17a
-log10(p.adj)
Expression (Log2 Fold Change)
p-adj < 0.05 and L2FC < -log2(1.5)
p-adj < 0.05 and L2FC > log2(1.5)
p-adj >= 0.05 or |L2FC| <= log2(1.5)
D WT cells: +p38i vs w/o p38i
Upregulated by p38i
Downregulated by p38i
Gzmb
Prf1
Ifng
GTKO cells: +p38i vs w/o p38i
Serpina3g
Serpina3f
Rorc
E OT-I T cells
F Polyclonal CD8+ T cells
Control
Activated p38i +
Activated p38i -
WT
GTKO
GZMB eFluor660
IFNG AF700
CD25 BV605
SSC-A

**Extended View Figure 5. Analysis of signaling pathways in ABIN1-deficient T cells. (A-D)** The same RNAseq experiment as presented in Figure 5E-F. n = 3 biological replicates. (A) Principal component analysis of the samples. **(B)** Comparison of these experiments and the previous set of experiments (presented in Figure 4C-D) by plotting the fold changes of activated *Abin1*$^{GTKO/GTKO}$ OT-I *Rag2*$^{KO/KO}$ (GTKO) vs *Abin1*$^{WT/WT}$ OT-I *Rag2*$^{KO/KO}$ (WT) T cells in both sets of experiments. **(C)** A volcano plot showing expression changes of activated T cells treated p38 MAPK inhibitor (12.5 µM) for combined WT and GTKO samples. **(D)** A volcano plot showing expression changes of activated T cells treated or non-treated with p38 MAPK inhibitor (12.5 µM) for WT and GTKO samples separately. **(E-F)** Lymph node cells from WT and GTKO **(E)** or polyclonal WT and GTKO **(F)** mice from the experiment shown in Figure 5G-H. Representative dot plots showing the expression of indicated markers are shown. (E) n = 7 (WT), or 8 (GTKO) mice in 3 independent experiments, (F) n = 10 mice in 3 independent experiments.

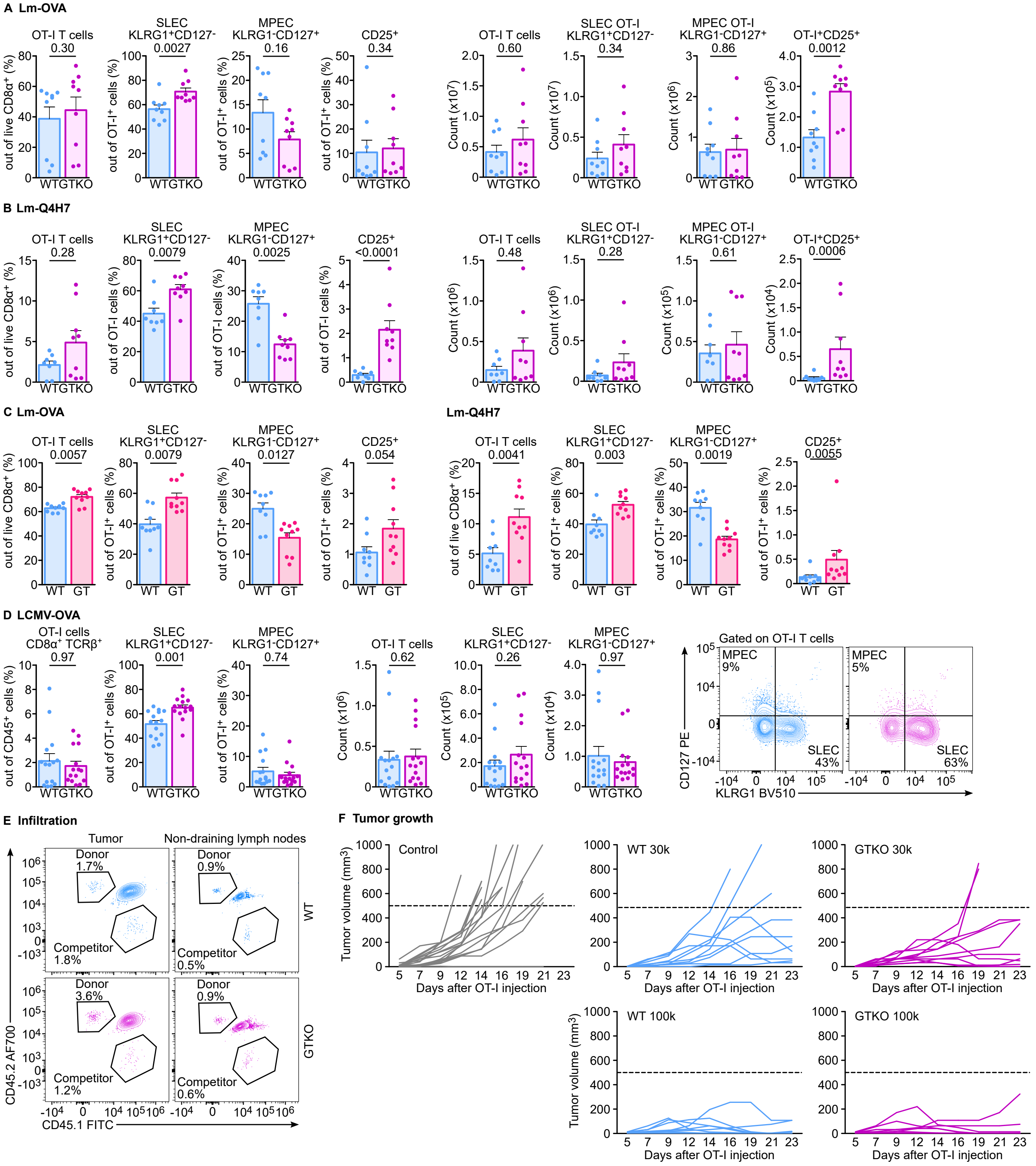
A Lm-OVA
OT-I T cells
0.30
out of live CD8α+ (%)
SLEC KLRG1+CD127-
0.0027
out of OT-I+ cells (%)
MPEC KLRG1-CD127+
0.16
CD25+
0.34
0.60
Count (x10^7)
SLEC OT-I KLRG1+CD127-
0.34
MPEC OT-I KLRG1-CD127+
0.86
Count (x10^6)
OT-I+CD25+
0.0012
Count (x10^5)
WT GTKO
B Lm-Q4H7
OT-I T cells
0.28
SLEC KLRG1+CD127-
0.0079
out of OT-I cells (%)
MPEC KLRG1-CD127+
0.0025
CD25+
<0.0001
0.48
Count (x10^6)
0.28
0.61
Count (x10^5)
OT-I+CD25+
0.0006
Count (x10^4)
C Lm-OVA
OT-I T cells
0.0057
SLEC KLRG1+CD127-
0.0079
MPEC KLRG1-CD127+
0.0127
CD25+
0.054
WT GT
Lm-Q4H7
0.0041
0.003
0.0019
0.0055
D LCMV-OVA
OT-I cells CD8α+ TCRβ+
0.97
out of CD45+ cells (%)
SLEC KLRG1+CD127-
0.001
MPEC KLRG1-CD127+
0.74
OT-I T cells
0.62
0.26
Count (x10^5)
0.97
Count (x10^4)
Gated on OT-I T cells
MPEC 9%
SLEC 43%
MPEC 5%
SLEC 63%
CD127 PE
KLRG1 BV510
E Infiltration
Tumor
Non-draining lymph nodes
Donor 1.7%
Competitor 1.8%
Donor 0.9%
Competitor 0.5%
WT
Donor 3.6%
Competitor 1.2%
Donor 0.9%
Competitor 0.6%
GTKO
CD45.2 AF700
CD45.1 FITC
F Tumor growth
Control
WT 30k
GTKO 30k
WT 100k
GTKO 100k
Tumor volume (mm³)
Days after OT-I injection

**Extended View Figure 6. (A-B)** The experiments as shown in Figure 6A-B with identical protocol, but using a transfer of $0.5\times10^5$ T cells. (A) n = 9 mice per group in 3 independent experiments. (B) n = 8 (WT), or 9 (GTKO) mice in 3 independent experiments. Quantified frequencies and counts of indicated subsets of donor cells are shown. **(C)** $1\times10^5$ OT-I cells from WT or *Abin1*$^{GT/GT}$ OT-I *Rag2*$^{KO/KO}$ (GT) mice were adoptively transferred to Ly5.2 hosts that were infected with Lm-OVA. Splenocytes were analyzed by flow cytometry on day 6 post infection. The frequency of indicated subsets are shown. n = 9 (WT) or 10 (GT) mice in 3 independent experiments. **(D)** $1\times10^4$ OT-I cells from WT or *Abin1*$^{GTKO/GTKO}$ OT-I *Rag2*$^{KO/KO}$ mice were adoptively transferred to the CD3ε$^{-/-}$ hosts that were infected with LCMV expressing OVA peptide. Splenocytes were analyzed by flow cytometry on day 5 post infection. Representative dot plots and quantified frequencies and absolute counts of indicated subsets of donor cells are shown. n = 15 mice per group in 3 independent experiments. **(E)** The experiment shown in Figure 6C. A representative experiment for tumor and non-draining lymph nodes. **(F)** The experiment shown in Figure 6D. The tumor growth in individual mice is shown. The dashed line represents the endpoint of the experiment (tumor volume 500 mm$^3$). The genotype and number of transferred OT-I *Rag2*$^{KO/KO}$ T cells are indicated.

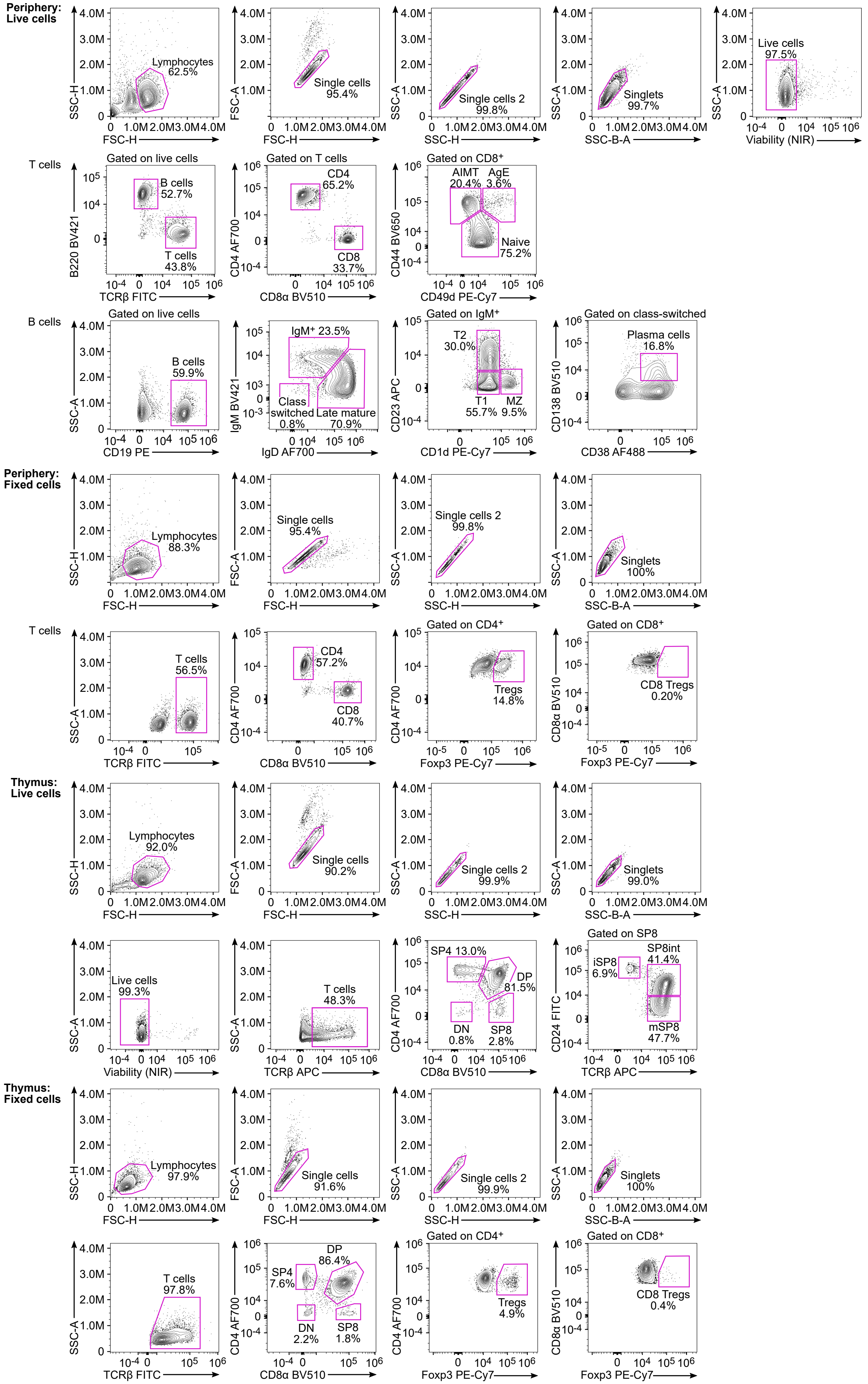

Periphery: Live cells
Lymphocytes 62.5%
Single cells 95.4%
Single cells 2 99.8%
Singlets 99.7%
Live cells 97.5%
SSC-H
FSC-H
FSC-A
SSC-A
SSC-B-A
Viability (NIR)
T cells
Gated on live cells
B cells 52.7%
T cells 43.8%
B220 BV421
TCRβ FITC
Gated on T cells
CD4 65.2%
CD8 33.7%
CD4 AF700
CD8α BV510
Gated on CD8+
AIMT 20.4%
AgE 3.6%
Naive 75.2%
CD44 BV650
CD49d PE-Cy7
B cells
Gated on live cells
B cells 59.9%
CD19 PE
IgM+ 23.5%
Class switched 0.8%
Late mature 70.9%
IgM BV421
IgD AF700
Gated on IgM+
T2 30.0%
T1 55.7%
MZ 9.5%
CD23 APC
CD1d PE-Cy7
Gated on class-switched
Plasma cells 16.8%
CD138 BV510
CD38 AF488
Periphery: Fixed cells
Lymphocytes 88.3%
Single cells 95.4%
Single cells 2 99.8%
Singlets 100%
T cells
T cells 56.5%
CD4 57.2%
CD8 40.7%
Gated on CD4+
Tregs 14.8%
Foxp3 PE-Cy7
Gated on CD8+
CD8 Tregs 0.20%
Thymus: Live cells
Lymphocytes 92.0%
Single cells 90.2%
Single cells 2 99.9%
Singlets 99.0%
Live cells 99.3%
T cells 48.3%
TCRβ APC
SP4 13.0%
DP 81.5%
DN 0.8%
SP8 2.8%
Gated on SP8
iSP8 6.9%
SP8int 41.4%
mSP8 47.7%
CD24 FITC
Thymus: Fixed cells
Lymphocytes 97.9%
Single cells 91.6%
Single cells 2 99.9%
Singlets 100%
T cells 97.8%
SP4 7.6%
DP 86.4%
DN 2.2%
SP8 1.8%
Gated on CD4+
Tregs 4.9%
Gated on CD8+
CD8 Tregs 0.4%

**Appendix 1.** Gating strategies for the analysis of live and fixed peripheral lymphocytes and thymocytes by flow cytometry used throughout this article.

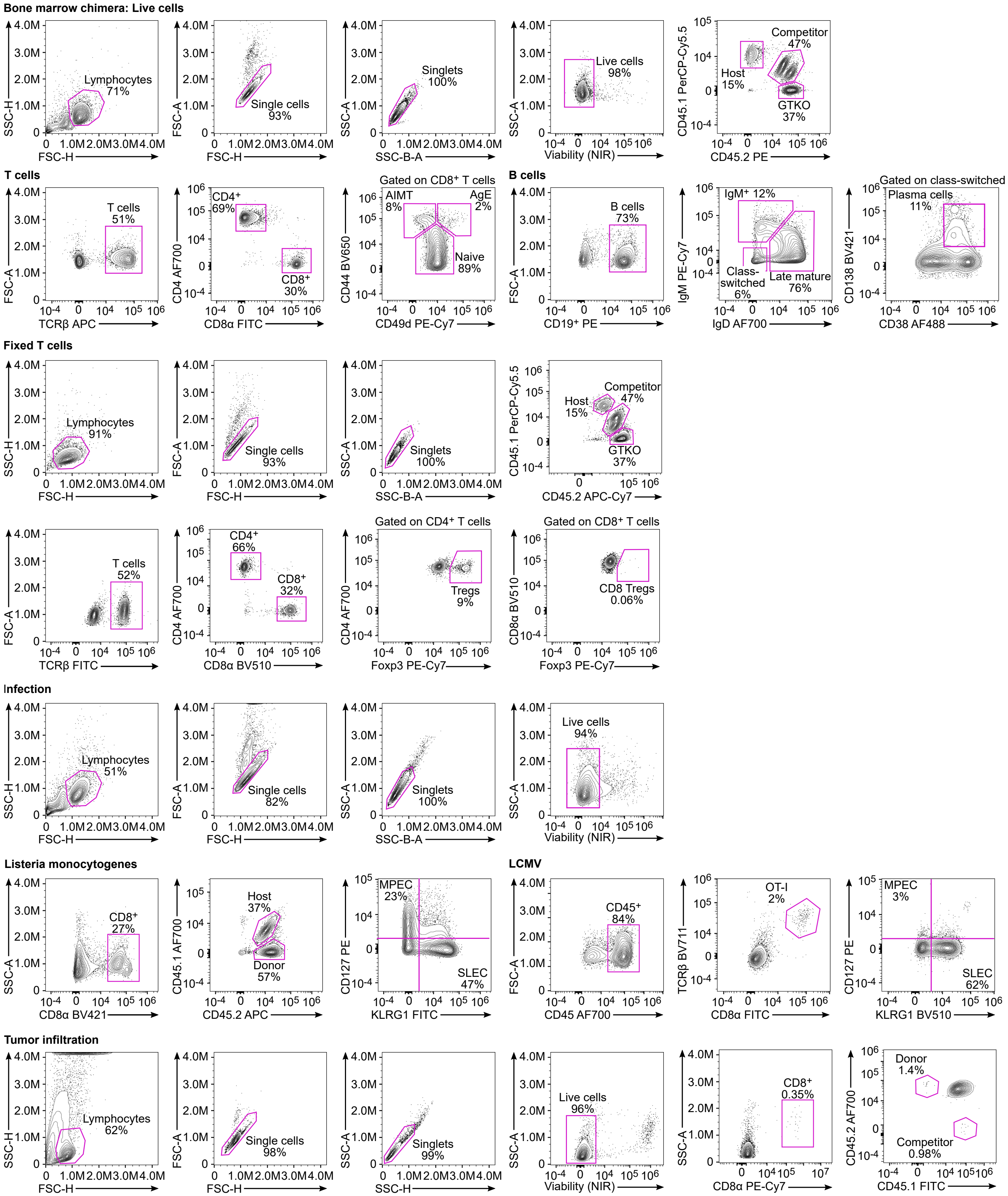
Bone marrow chimera: Live cells
Lymphocytes 71%
Single cells 93%
Singlets 100%
Live cells 98%
Competitor 47%
Host 15%
GTKO 37%
SSC-H
FSC-H
FSC-A
SSC-A
SSC-B-A
Viability (NIR)
CD45.1 PerCP-Cy5.5
CD45.2 PE
T cells
T cells 51%
TCRβ APC
CD4+ 69%
CD8+ 30%
CD4 AF700
CD8α FITC
Gated on CD8+ T cells
AIMT 8%
AgE 2%
Naive 89%
CD44 BV650
CD49d PE-Cy7
B cells
B cells 73%
CD19+ PE
IgM+ 12%
Class-switched 6%
Late mature 76%
IgM PE-Cy7
IgD AF700
Gated on class-switched
Plasma cells 11%
CD138 BV421
CD38 AF488
Fixed T cells
Lymphocytes 91%
Single cells 93%
Singlets 100%
Host 15%
Competitor 47%
GTKO 37%
CD45.2 APC-Cy7
T cells 52%
TCRβ FITC
CD4+ 66%
CD8+ 32%
CD8α BV510
Gated on CD4+ T cells
Tregs 9%
Foxp3 PE-Cy7
Gated on CD8+ T cells
CD8 Tregs 0.06%
CD8α BV510
Infection
Lymphocytes 51%
Single cells 82%
Singlets 100%
Live cells 94%
Listeria monocytogenes
CD8+ 27%
CD8α BV421
Host 37%
Donor 57%
CD45.1 AF700
CD45.2 APC
MPEC 23%
SLEC 47%
CD127 PE
KLRG1 FITC
LCMV
CD45+ 84%
CD45 AF700
OT-I 2%
TCRβ BV711
CD8α FITC
MPEC 3%
SLEC 62%
KLRG1 BV510
Tumor infiltration
Lymphocytes 62%
Single cells 98%
Singlets 99%
SSC-H
Live cells 96%
CD8+ 0.35%
CD8α PE-Cy7
Donor 1.4%
Competitor 0.98%
CD45.2 AF700
CD45.1 FITC

**Appendix 2.** Gating strategy used in analysis of bone marrow chimeras, infection *(Listeria* and LCMV), and cell infiltration in tumor by flow cytometry used in this article.

**Immunophenotyping of *Abin1*$^{dE5/dE5}$ mice**

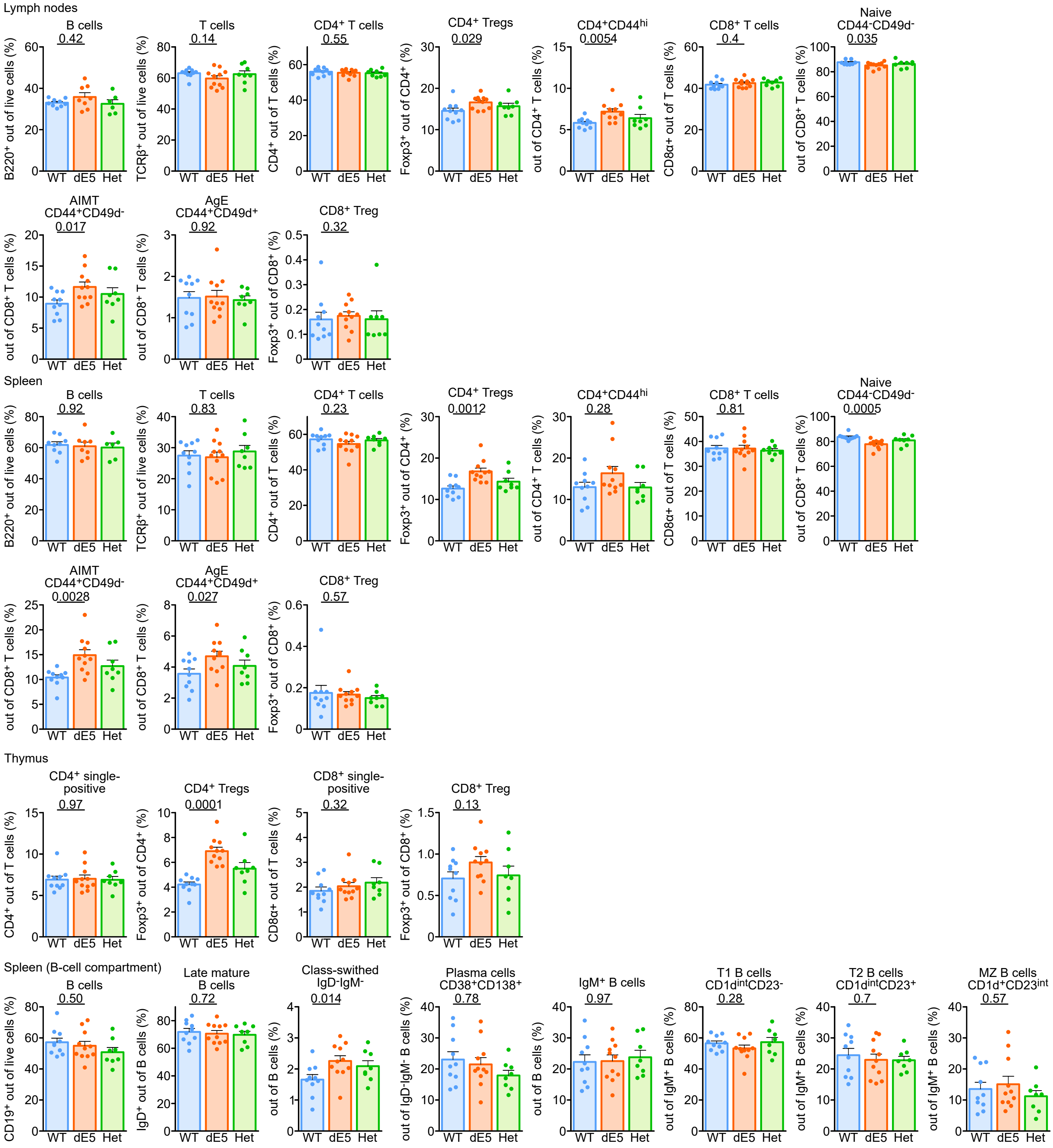

**Appendix 3.** Immunophenotyping of *Abin1*$^{dE5/dE5}$ mice by flow cytometry. Quantification of indicated subsets are shown. The results are shown as means + SEM and p-values are indicated. Statistical significance was determined by two-tailed Mann-Whitney test.

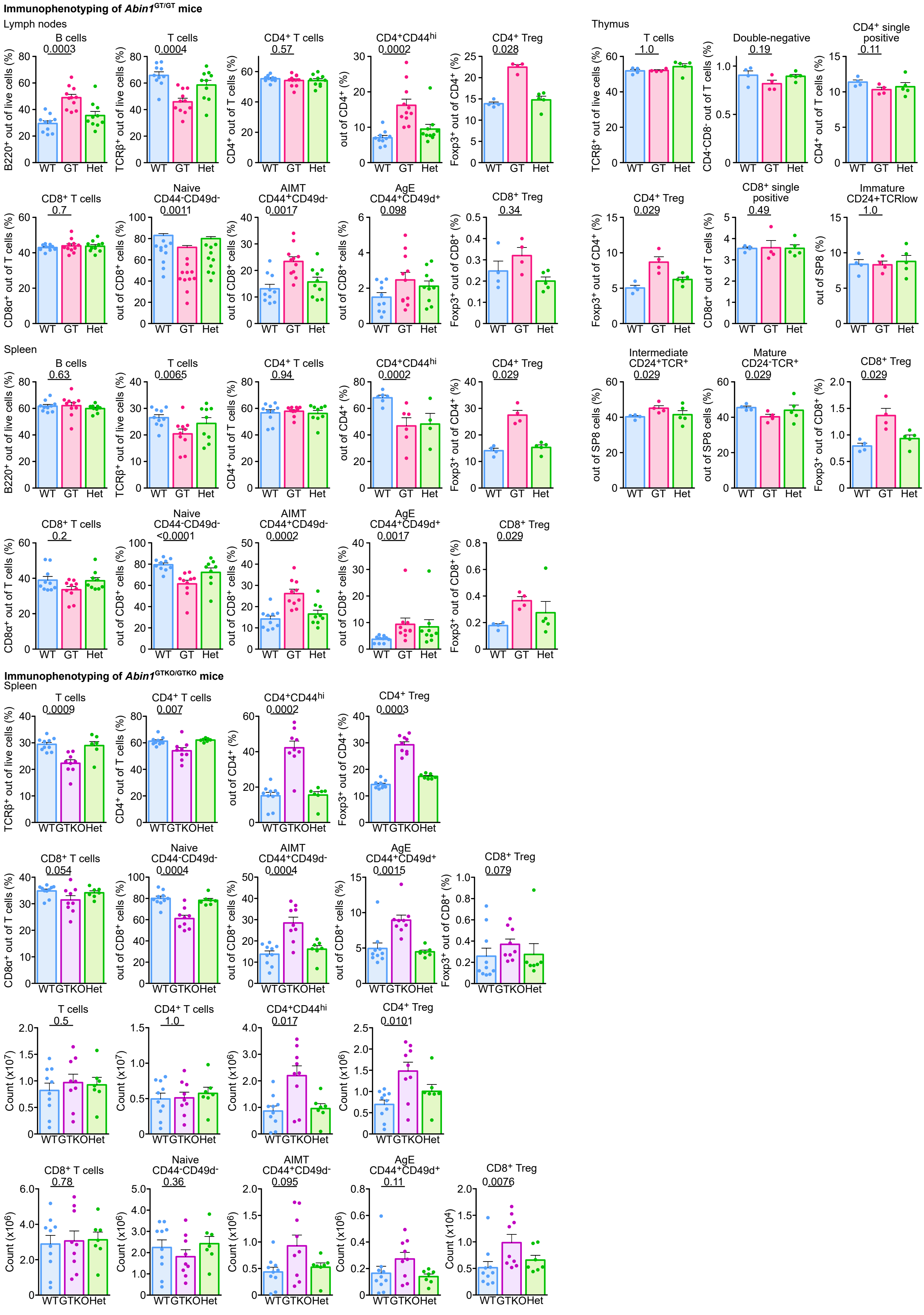
Immunophenotyping of Abin1GT/GT mice
Lymph nodes
B cells 0.0003
T cells 0.0004
CD4+ T cells 0.57
CD4+CD44hi 0.0002
CD4+ Treg 0.028
CD8+ T cells 0.7
Naive CD44-CD49d- 0.0011
AIMT CD44+CD49d- 0.0017
AgE CD44+CD49d+ 0.098
CD8+ Treg 0.34
Thymus
T cells 1.0
Double-negative 0.19
CD4+ single positive 0.11
CD4+ Treg 0.029
CD8+ single positive 0.49
Immature CD24+TCRlow 1.0
Intermediate CD24+TCR+ 0.029
Mature CD24-TCR+ 0.029
CD8+ Treg 0.029
Spleen
B cells 0.63
T cells 0.0065
CD4+ T cells 0.94
CD4+CD44hi 0.0002
CD4+ Treg 0.029
CD8+ T cells 0.2
Naive CD44-CD49d- <0.0001
AIMT CD44+CD49d- 0.0002
AgE CD44+CD49d+ 0.0017
CD8+ Treg 0.029
WT GT Het
Immunophenotyping of Abin1GTKO/GTKO mice
Spleen
T cells 0.0009
CD4+ T cells 0.007
CD4+CD44hi 0.0002
CD4+ Treg 0.0003
CD8+ T cells 0.054
Naive CD44-CD49d- 0.0004
AIMT CD44+CD49d- 0.0004
AgE CD44+CD49d+ 0.0015
CD8+ Treg 0.079
T cells 0.5
CD4+ T cells 1.0
CD4+CD44hi 0.017
CD4+ Treg 0.0101
CD8+ T cells 0.78
Naive CD44-CD49d- 0.36
AIMT CD44+CD49d- 0.095
AgE CD44+CD49d+ 0.11
CD8+ Treg 0.0076
WT GTKO Het

**Appendix 4.** Immunophenotyping of *Abin1*$^{GT/GT}$ and *Abin1*$^{GTKO/GTKO}$ mice by flow cytometry. Quantifications of indicated subsets are shown. The results are shown as means + SEM and p-values are indicated. Statistical significance was determined by two-tailed Mann-Whitney test.

**Supplemental Table 1.**

Normalized counts generated by DESeq2 for bulk RNAseq data (Figure EV4B) of *Abin1*$^{GTKO/GTKO}$ and WT OT-I T cells activated or not with anti-CD3/CD28 beads. The last 3 columns denote log2-fold changes, p-values, and adjusted p-values for contrast between the cells of *Abin1*$^{GTKO/GTKO}$ vs. WT OT-I T cells (irrespective of activation).

**Supplemental Table 2.**

Normalized counts generated by DESeq2 for bulk RNAseq data (Figure EV5A) of *Abin1*$^{GTKO/GTKO}$ and WT OT-I T cells activated with anti-CD3/CD28 beads and treated or not with the p38 inhibitor. Log2-fold changes, p-values, and adjusted p-values for the contrast between cells of *Abin1*$^{GTKO/GTKO}$ vs. activated WT OT-I T cells (irrespective of p38 inhibition) and the contrast between T cells without vs. with p38 inhibitor (irrespective of the genotype) are shown.

**Supplemental Table 3.**

The list of genes up-regulated (fold change > 1.5) in activated *Abin1*$^{GTKO/GTKO}$ vs. activated WT OT-I T cells in both experiments (i.e., Figure EV4B and EV5A) used for the GSEA (Figure 4E).
